# Landscape of immune-related signatures induced by targeting of different epigenetic regulators in melanoma: implications for immunotherapy

**DOI:** 10.1101/2022.04.13.488140

**Authors:** Andrea Anichini, Alessandra Molla, Gabriella Nicolini, Valentina E. Perotti, Francesco Sgambelluri, Alessia Covre, Carolina Fazio, Maria Fortunata Lofiego, Anna Maria di Giacomo, Sandra Coral, Antonella Manca, Maria Cristina Sini, Marina Pisano, Teresa Noviello, Francesca Caruso, Silvia Brich, Giancarlo Pruneri, Andrea Maurichi, Mario Santinami, Michele Ceccarelli, Giuseppe Palmieri, Michele Maio, Roberta Mortarini, the EPigenetic Immune-oncology Consortium AIRC (EPICA) investigators

## Abstract

**Background:** Innovative cancer immunotherapy approaches aim at combining immune checkpoint inhibitors with other immunomodulatory agents. Epigenetic regulators can control immune-related genes, therefore targeting them with specific inhibitors may be a potential way forward. Here we identified immune-related signatures induced by four classes of epigenetic drugs in human melanoma cells to define the most promising agent and to understand its biological activity in-vitro, in-vivo and in clinical samples.

**Methods:** Human melanoma cell lines were characterized for mutational and differentiation profile and treated with inhibitors of DNA methyltransferases (guadecitabine), histone deacetylases (givinostat), bromodomain and extraterminal domain proteins (JQ1 and OTX-015) and enhancer of zeste homolog 2 (GSK126). Drug-specific gene signatures were identified by Clariom S and Nanostring platforms. Modulation of 14 proteins was determined by quantitative western blot. Ingenuity Pathway Analysis (IPA) identified Upstream Regulator (UR) molecules explaining changes in gene expression and biological activity of drugs. Gene set enrichment and IPA were used to test modulation of guadecitabine-specific gene and UR signatures, respectively, in on-treatment tumor biopsies from melanoma patients enrolled in the Phase Ib NIBIT-M4 Guadecitabine + Ipilimumab Trial.

**Results:** Drug-specific gene and UR signatures were identified for each of the four inhibitors. Immune-related genes were frequently upregulated by guadecitabine, to a lesser extent by givinostat, but downregulated by JQ1 and OTX-015. GSK126 was the least active drug. Treatment of melanoma cells with combination of two epigenetic drugs revealed a dominant effect of guadecitabine and JQ1 on immune-related gene modulation. Drug-specific modulatory profiles were confirmed at the protein level. The guadecitabine-specific UR signature was characterized by activated molecules of the TLR, NF-kB, and IFN innate immunity pathways and was induced in drug-treated melanoma, mesothelioma, hepatocarcinoma cell lines and human melanoma xenografts. Most of the guadecitabine-specific signature genes (n>160) were upregulated in on-treatment tumor biopsies from NIBIT-M4 trial. Progressive activation of guadecitabine UR signature molecules was observed in on-treatment tumor biopsies from responding compared to non-responding patients.

**Conclusions:** Guadecitabine was the most promising immunomodulatory agent among those investigated. This DNA methyltransferases inhibitor emerged as a strong inducer of innate immunity pathways, supporting the rationale for its use in combinatorial immunotherapy approaches.

## Introduction

Following the approval 10 years ago of the anti-CTLA-4 antibody ipilimumab for treatment of metastatic melanoma [1], the management of an increasing spectrum of solid tumors has been revolutionized by the introduction of immunotherapy based on immune checkpoint inhibitors (ICI). Antibodies targeting CTLA-4 or the PD-1/PD-L1 axis have been approved for the treatment of 19 cancer types and two tissue-agnostic conditions [2]. A recent meta-analysis on 23,760 patients and 37 Phase II/III trials, has shown that ICI immunotherapy provides a survival advantage compared to control treatments irrespective of age, sex and ECOG performance status [3]. In spite of these remarkable clinical achievements, only a fraction of patients obtains durable complete responses, even in the most responsive tumor types, and both intrinsic and acquired resistance to ICI is frequent [4]. Therefore, there is an urgent need to design improved immunotherapy approaches. One of the options being tested in early Phase I/II trials is the association of ICIs with immunomodulatory drugs [5,6]. In support of these combinatorial immunotherapy approaches, the recently completed Italian Network for Tumor Biotherapy (NIBIT) Phase Ib NIBIT-M4 trial [6], based on the association of ipilimumab with guadecitabine in advanced melanoma, has shown significant tumor immunomodulatory effects and preliminary evidence of promising clinical activity.

Inhibitors targeting molecules involved in epigenetic regulation have attracted considerable interest in the field of immuno-oncology, as several of these drugs have immune-related effects on tumor cells, as well as on immune cells, that could potentially synergize with ICI [5, 7, 8]. A large series of preclinical studies has shown that epigenetic drugs targeting DNA methyl transferases (DNMT), histone deacetylases (HDAC), enhancer of zeste homolog 2 (EZH2), bromodomain and extra-terminal domain (BET) proteins, and lysine-specific demethylase 1 (LSD-1), when combined with ICI, can contribute to the overall anti-tumor efficacy of immunotherapy, by affecting the structure of the tumor immune landscape, the generation of anti-tumor immunity, and by modulating immunosuppressive mechanisms and immune escape processes [5, 7, 8]. On the other hand, neither single epigenetic drugs, nor any class of drugs targeting epigenetic regulators, can elicit all of these immunomodulatory effects. Instead, there is a prominent drug-related specificity. For example, DNMT inhibitors can prevent T cell exhaustion [9], activate the interferon response pathway [10], rescue cGAS and STING gene expression [11] and improve antigen processing and presentation [12]. HDAC inhibitors, alone or in association with DNMT inhibitors, can promote T cell recruitment at tumor site [13, 14], reprogram tumor associated macrophages and reduce frequency/suppressive function of Tregs and MDSCs [15, 16]. EZH2 inhibitors can rescue MHC class I transcription [17] and counteract melanoma dedifferentiation [18]. BET inhibitors can downregulate PD-L1 expression [19] and maintain CD8^+^ T stem cell memory and T central memory differentiation states [20].

The emerging picture is a heterogeneous landscape of diverse and sometimes opposite immune-related activities on either tumors or the tumor microenvironment, or both, by different classes of epigenetic drugs [21, 22]. This immunomodulatory complexity contributes to explain why early combination immunotherapy trials are not focusing on a single epigenetic drug or a drug class. In fact, among >80 recently reviewed Phase I/II combination immunotherapy trials, 15 different inhibitors directed at five classes of epigenetic targets are being tested in association with 9 different ICI [5].

To foster progress in the field of epigenetic immuno-modulation here we carried out a comparative profiling of signatures induced by four epigenetic drugs representing the classes of DNMT, HDAC, BET and EZH2 inhibitors and tested in the melanoma context, a tumor type currently treated by immunotherapy. We defined the breadth of drug-specific immune-related signatures in neoplastic cells fully characterized for genetic background and differentiation profile. We found that guadecitabine is a strong immunomodulatory agent promoting activation of innate immunity pathways both in-vitro and in-vivo in patients treated with this DNMT inhibitor.

## Materials and Methods

### Melanoma cell lines

Melanoma cell lines (n=14) were established, maintained and routinely tested for the absence of mycoplasma contamination by PCR, as previously described [23]. Origin of the cell lines from primary lesions (n=1) or from metastases (n=13), histopathological features of the corresponding primary melanoma, cell line authentication by STR profiling (Gene-Print10 kit, Promega) and mutational profile by targeted NGS are described in Table S1A-D. The melanoma cell lines were representative of the main molecular subtypes: BRAF^mut^/NRAS^wt^ (n=10), BRAF^wt^/NRAS^mut^ (n=3), and BRAF^wt^/ NRAS^wt^ (n=1). Expression in n=10 melanoma cell lines of the genes targeted by epigenetic drugs (Fig. S1) was evaluated as described in Supplementary Methods. Mesothelioma cell lines (MPP-89, MM98 and MES2a) were obtained from pleural effusion of mesothelioma patients, cultured, and treated with guadecitabine as previously described [24].

### Epigenetic drugs and treatments

Melanoma cell lines were treated with the following drugs: the DNMT inhibitors Decitabine (Selleckchem) and Guadecitabine (MedChemExpress LLC), the HDAC inhibitor Givinostat (ITF-2357, Selleckchem), the BET inhibitors JQ-1 (Selleckchem) and OTX015 (Selleckchem), the EZH2 inhibitor GSK-126 (Selleckchem), and the CDK4/6 inhibitor Abemaciclib (Selleckchem). Susceptibility of ten melanoma cell lines to a range of doses between 7.8 and 3,000 nM of all the drugs was assessed by the 3-(4,5)dimethylthiazol-2,5-diphenyltetrazolium bromide (MTT) assay at 96h as described [25]. On the basis of the dose-response curves, drug doses for all subsequent experiments were tailored for each cell line and for each drug to achieve, in all instances, a response >60% of control in the MTT assay (Table S2A, B). For all subsequent assays, melanoma cells were seeded at 1.25×10^4^/mL in T75 flasks (Greiner Bio-One) with RPMI-1640 medium (Life Technologies Limited) supplemented with 4% FCS (Biological Industries) without antibiotics. Since the activity of epigenetic drugs is coupled to cell proliferation [26], melanoma cells were treated twice with each drug (at T=24h and T=72h) and then evaluated at T=144h. For some experiments melanoma cells were treated with all possible combinations of two out of the four epigenetic drugs.

### NGS analysis

DNA was extracted from 0.5-1×10^6^ melanoma cell lines by Purelink Genomic DNA Mini Kit (Invitrogen). RNAse treatment was carried out by RNAse cocktail (Invitrogen). Quantitative assessment of DNA was performed by a Nanodrop 2000 spectrophotometer and DNA integrity was confirmed by gel electrophoresis. Next generation sequencing (NGS) assays on cell lines DNA were done as described in Supplementary Methods.

### RNA extraction and gene expression analysis

RNA was extracted from melanoma cell lines by TRIzol (Thermo Fisher Scientific). Melanin removal was carried out by the CTAB-UREA method [27]. Samples were treated with DNase (Qiagen) and purified by RNeasy MinElute Cleanup Kit (Qiagen). The quality of total RNA was first assessed using the RNA 6000 Pico Assay RNA chips run on an Agilent Bioanalyzer 2100 (Agilent Technologies). RNA was also extracted from tumor nodules previously obtained from immunodeficient mice treated or nor with guadecitabine and bearing human melanoma cell line 195 xenografts [28, 29]. Gene expression analysis was performed as described in Supplementary Methods.

### Quantitative Western blot

SDS-PAGE was performed with 30 μg of protein lysate on 4– 12% NuPAGE Bis-Tris (Thermo Fisher Scientific), in MOPS buffer as described [23]. Primary antibodies (Table S3) were diluted in milk 5% or BSA 5% in TBST as described [23] and incubated overnight. Development was performed with the ECL normal western blot detection system by the chemiluminescence method. Images were acquired with the Alliance Imaging System (Uvitec). Densitometric analysis was carried out by Quantity One software (Bio-Rad Laboratories Inc.). Normalized treated/control ratios were computed on the basis of background-adjusted density values and then visualized by a color code.

### Data analysis

The Transcriptomics Analysis Console (TAC) software (Applied Biosystems) was used to identify significantly modulated genes by treatment of two melanoma cell lines (VRG100 and CST30) treated with Guadecitabine, Givinostat, JQ1, OTX-015, GSK126 or Abemaciclib, and of melanoma nodules removed from immunodeficient mice treated with Guadecitabine. Analysis settings were as follows: gene level fold change >|1.2|, gene-level p value: <0.05; gene-level FDR: <0.05. Nanostring data were analyzed with nSolver software 4.0 (Nanostring Technologies) as described in Supplementary Methods. VENNTURE software [30], allowing Edwards-Venn diagram generation for multiple dataset analysis, was used to classify all genes with significant modulation by any of the treatments. Ingenuity Pathway Analysis (IPA 8.5, www.ingenuity.com) was used, as described in Supplementary Methods, to carry out Upstream Regulator (UR) analysis on: a) significantly modulated genes by different treatments; b) differentially expressed genes in pre-therapy and on-treatment lesions from responding vs non responding patients from the NIBIT-M4 trial [6]. IPA was also used for canonical pathway analysis as described in Supplementary Methods. RNA-seq and DNA methylation data from tumor samples obtained at baseline (week 0), at week 4 and week 12 of treatment were retrieved from the published NIBIT M4 Phase Ib trial [6]. Raw data as FASTQ files of melanoma patient’s biopsies were aligned to the human reference genome (GRCh38/hg38) using STAR version 2.7.0b [31] and the gene expression level was quantified using featureCounts version 1.6.3 [32]. Downstream analyses were performed in the R statistical environment. EDASeq package [33] was used to normalize the count matrix using GC-correction for the within normalization step and upper-quantile for the between phase. EdgeR package [34] was used to identify differentially expressed genes at week4 and week12 compared to week0 and at week12 compared to week4. The guadecitabine-specific gene signature (upregulated genes), identified in this manuscript, was used to construct a heatmap of the Log2 fold changes for the three different comparisons. ClusterProfiler package [35] was used to perform and visualize GSEA pre-ranked analyses using the guadecitabine-specific gene signature in each comparison.

## Results

### Epigenetic drugs modulate expression of 20 gene families

A panel of melanoma cell lines was fully characterized for tissue of origin (Table S1A), expression of epigenetic drug targets (Fig. S1), mutational profile (Table S1D), differentiation-related signatures (Fig. S2) as defined by Tsoi et al [36], as well as for susceptibility to four epigenetic drugs (Fig. S3 and Table S2A, B). Two of these cell lines (VRG100 and CST30) were selected as examples of divergent transcriptomic profiles found in melanoma (Fig. S4). Gene modulation in these two lines by DNMT (decitabine/guadecitabine), HDAC (givinostat), BET (JQ1) and EZH2 (GSK126) inhibitors was investigated by whole genome gene expression analysis. The CDK4/6 inhibitor abemaciclib, a drug with immunomodulatory activity unrelated to epigenetic regulation [37], was also included in these experiments. The five drugs modulated 6% to >30% of the genes in the two cell lines (Fig. S5) and, based on Edwards-Venn diagrams [30], showed both drug-specific as well as shared modulatory activities (Fig. S6). Guadecitabine, givinostat, JQ1, GSK-126 and abemaciclib exerted pleiotropic modulatory activities on 20 gene families grouped into 7 biological functions (Fig. 1), including epigenetic regulation, proliferation, immune regulation, enzymatic activity, structural function, adhesion and cellular differentiation. The drugs elicited variable modulatory activity on epigenetic regulator genes (DNMT, HDAC, BET and PRC2 components), and on genes encoding histones and nuclear pore complex interacting proteins (NPIPA genes). Guadecitabine, givinostat and JQ1 were the most active drugs, followed by GSK-126, in modulation of genes encoding HLA-I/II and antigen processing machinery (APM) components, heat shock proteins, IFN (IFI and IRF), TNF/TNFR and TGFβ pathways. These immune-related genes were frequently upregulated by guadecitabine and givinostat, but downregulated by JQ1 (Fig. 1).

**Figure 1.**
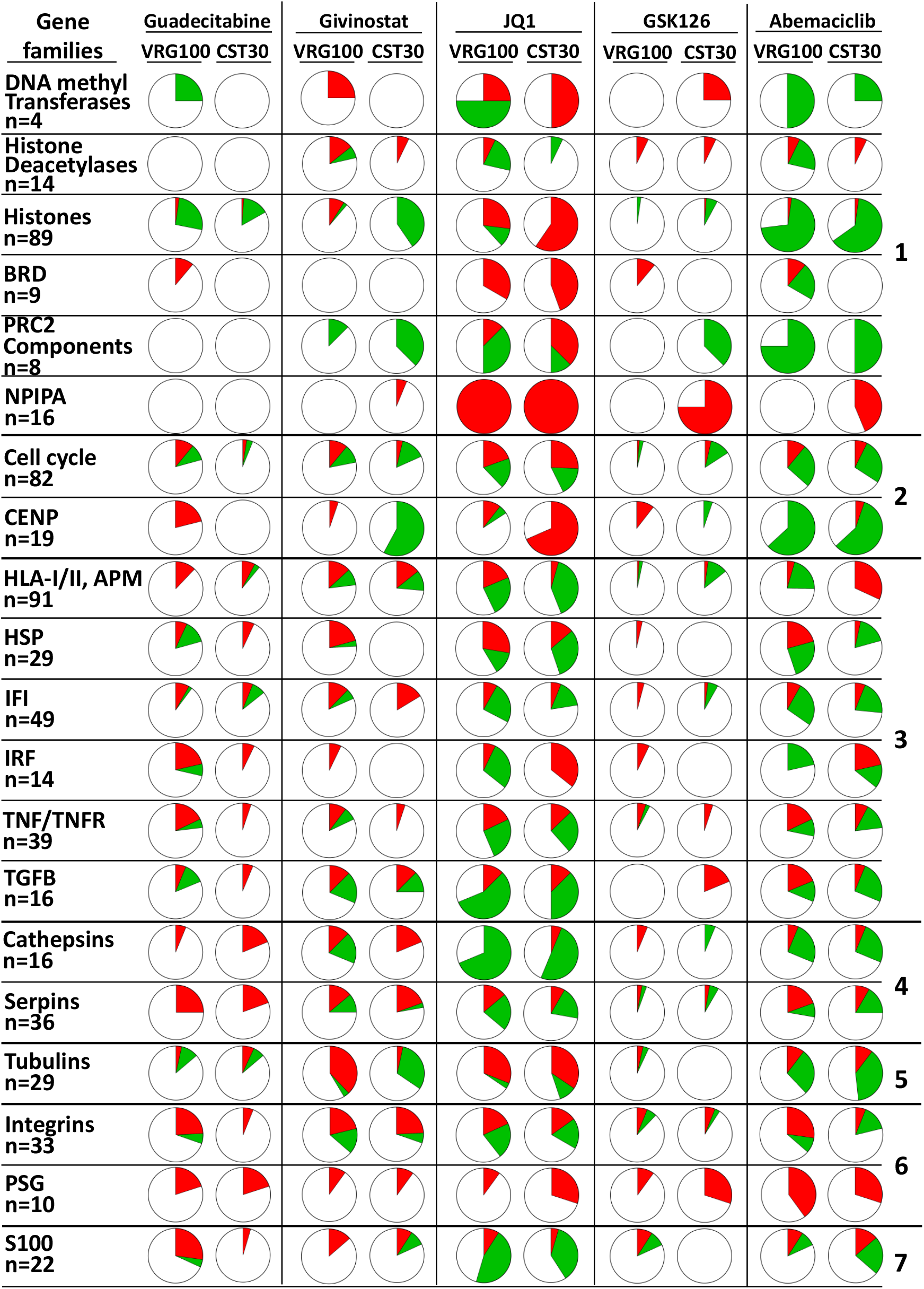
Landscape of gene modulation by epigenetic drugs in melanoma cell lines. Each pie chart shows the % of genes significantly modulated in each of 20 families or functional groups (red=upregulated genes, green=downregulated genes) by each drug in the two cell lines (VRG100 and CST30), on the basis of FC and p value. The 20 gene families are further classified into 7 groups identified on the right hand side of the figure: 1, epigenetic regulation; 2, proliferation; 3, immune regulation; 4, enzymes; 5, structural components; 6, adhesion; 7, differentiation.

### Guadecitabine is the most active drug in upregulation of immune-related genes, while JQ1 is a strong suppressor of immune-related gene expression

Immunomodulation emerged as a dominant feature of the epigenetic drugs in the two cell lines VRG100 and CST30. Therefore, we focused on the quantitative determination of 731 immune-related genes, present in the Nanostring Cancer Immune panel and re-classified by us into 21 functional classes (Fig. S7). In these experiments we used the whole panel of melanoma cell lines. Quantitative analysis (Fig. S8) and hierarchical clustering of Nanostring data, according to the 21 functional classes (Fig. S9), indicated that guadecitabine and givinostat had a predominantly positive effect on immune-related gene expression, while JQ1 induced frequent gene downmodulation. To reduce the complexity emerging from the gene-level analysis (shown in Table S4) of the modulatory effects of all drugs on the 21 functional classes, we used *ad-ho*c developed metrics. This approach (Fig. 2) allowed to visualize the proportion of genes being significantly modulated in each gene class and the direction of the most frequent effects by each drug (in terms of up-or down-regulated genes). Guadecitabine and givinostat mostly induced upregulation of immune-related genes in the majority of the cell lines, while the BET inhibitor JQ1 produced the opposite effects (Fig. 2). GSK126 and abemaciclib were the least active drugs and also showed cell line-specific effects. Immune-related gene modulation by the three most active drugs was observed across the melanoma cell line panel without any clear association with the mutational or differentiation profile of the cell lines (Fig. 2). Two of the cell lines (BNV13 and GRD43) were treated with decitabine, the active metabolite of guadecitabine, yielding similar results to the cell lines treated with guadecitabine (Fig. 2). Guadecitabine was the most active drug in upregulation of genes belonging to the following functional classes: cancer testis, adhesion molecules, positive/negative co-stimulation, myeloid-related, cytokines and receptors, T/NK related, immune cell lineage/differentiation, regulation of inflammation, TNF/TNFR pathway, Type I-II-III IFN pathways and intracellular signaling (Fig. 2 and Table S4 for gene level analysis). The BET inhibitor JQ1 was the most active drug in inducing downregulation of selected genes related to immune cells lineage/differentiation markers, TNF/TNFR pathway, Type I-II-III IFNs pathways, intracellular signaling and transcriptional regulation, TLR pathway, HLA Class I/II and antigen presentation pathways (Fig. 2 and Table S4 for gene-level analysis). Nanostring analysis of a subset of six cell lines treated with a different BET inhibitor (OTX-015), tested in clinical trials [38], confirmed the predominant down-modulation of immune-related genes by this class of epigenetic drugs (Fig. S10).

**Figure 2.**
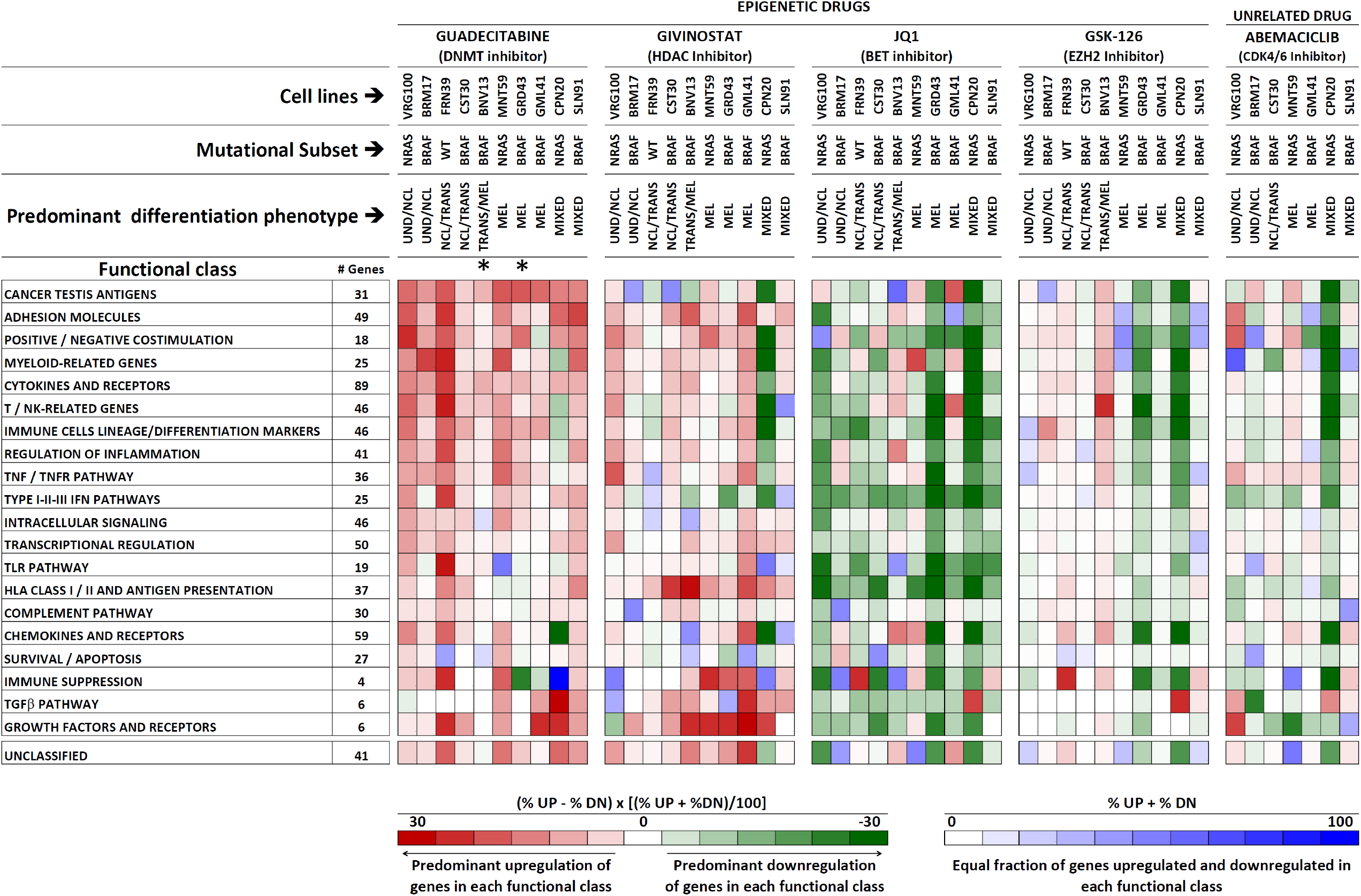
Immune-related gene modulation by epigenetic drugs in melanoma. Modulation of 731 genes was assessed by the Nanostring Cancer Immune panel upon treatment with 4 epigenetic drugs and with Abemaciclib of ten melanoma cell lines with the indicated mutational and differentiation profiles (as defined in Tables S1 and Fig. S2, respectively). Genes in the Nanostring panel were grouped into 21 functional classes (as described in Fig. S7). The overall modulatory activity of all drugs on each gene class was visualized by *ad-hoc* developed, color-coded metrics described at the bottom of the Figure. The first metric adopts a red/green color code to visualize instances of predominant up-or down-regulation of genes by each drug and in each cell line. This metric contains a normalization factor (=[(%UP+%DN)/100)]) that takes into account the different number of genes modulated in each cell line. The second metrics adopts shades of blue to visualize instances where an equal fraction of up-and down-modulated genes in each class was observed. Up-or down-modulation of each gene by any drug was defined on the basis of a Treated/Control ratio >|1.5|. *: these two cell lines were treated with decitabine, the active metabolite of guadecitabine.

By exploiting Nanostring data from all cell lines we derived all the epigenetic drug-specific gene signatures listing the most frequently modulated genes (Fig. S11). The guadecitabine-specific and the JQ1-specific gene signatures contained each ∼170 genes, mostly being upregulated by the former drug while being downregulated by the latter. The givinostat-specific gene signature contained 128 genes (several shared with the guadecitabine signature) while the GSK126 signature contained 40 genes (Fig S11).

### Combinatorial treatments with epigenetic drugs indicate a dominant effect of guadecitabine and JQ1 on immune-related gene modulation

Melanoma cells from the panel were treated with all six binary combinations of the four epigenetic drugs (guadecitabine + givinostat, guadecitabine + JQ1, guadecitabine + GSK126, givinostat + JQ1, givinostat + GSK126, JQ1 + GSK126). Gene modulation induced by the combinations was then measured by Nanostring Cancer Immune panel and compared to the effects of single drugs (Table S5). In most instances, combinatorial treatments did not enhance gene modulation compared to single treatments. Moreover, the effect of a “dominant” drug was evident. In fact, association of guadecitabine with givinostat or with GSK-126 replicated the pattern of immune-related gene upregulation exerted by guadecitabine alone (Table S5). Similarly, any association of JQ1 with the other drugs replicated the dominant suppressive effect on immune-related gene expression exerted by JQ1 alone (Table S5). These results argue against the possibility of achieving more effective modulation of immune-related genes by combining two epigenetic drugs belonging to different classes.

### Immune-related genes modulated by guadecitabine belong to the “low expression/high methylation” class

Guadecitabine is expected to induce upregulation of genes subjected to negative regulation by promoter methylation [39]. Therefore genes in the guadecitabine signature should be characterized by high levels of methylation and low expression. To test this hypothesis, we retrieved data on gene expression and DNA methylation of a large set of melanoma cell lines from the CellMinerCDB database [40]. We defined a gene expression/methylation classification in 4 subsets (a,b,c and d in Fig. S12A-C) by setting appropriate methylation and expression thresholds. Then this classification was extended to all the genes in the guadecitabine-specific signature (Fig. S13). We found that most of them were indeed characterized by high methylation and low expression in most cell lines, namely they belong to group b). We then retrieved gene-specific methylation and gene expression data previously obtained on tumor samples obtained at week 0 (baseline, w0), at week 4 (w4) and week 12 (w12) in the context of the guadecitabine + ipilimumab NIBIT-M4 clinical trial [6]. We focused on the subset of the cancer testis (CT) genes, that can be modulated by guadecitabine and that have tumor-restricted expression [41]. By adopting a scoring system described in Fig. S14 A, B, we found frequent increased expression of CT genes associated with reduction of their methylation level in tumor samples obtained at w4 and w12 of treatment, compared to w0 biopsies (Fig. S14C).

### Epigenetic drugs modulate expression of immune-related proteins in melanoma

The immuno-related activity of the epigenetic drugs was assessed by quantitative Western blot analysis (pipeline of data analysis and visualization described in Fig. S15 and Fig. S16) by looking at expression and modulation of 14 immune-related and differentiation-related proteins in 11 melanoma cell lines (Fig. 3). Guadecitabine, followed by givinostat and GSK-126, were the most active drugs in increasing expression of HLA class I and II-related proteins (HLA Class I heavy chain, B2M and HLA-DR), but guadecitabine was more active compared to givinostat and to GSK-126 on molecules belonging to the antigen processing and presentation pathway (TAP1, TAP2, LMP2/PSMB9, Fig. 3). In contrast, the two BET inhibitors induced, in most instances, a reduction of expression of these classes of proteins. Guadecitabine and the BET inhibitors confirmed their opposite effects on expression of IFN-pathway related proteins (OAS3 and IFITM1), on PD-L1 / CD274 and on CEACAM1. Interestingly, all the 5 epigenetic drugs had a predominant inhibitory effect on expression of melanoma differentiation-related proteins MITF and SOX-10 and of the transcription factor MYC (Fig. 3).

**Figure 3.**
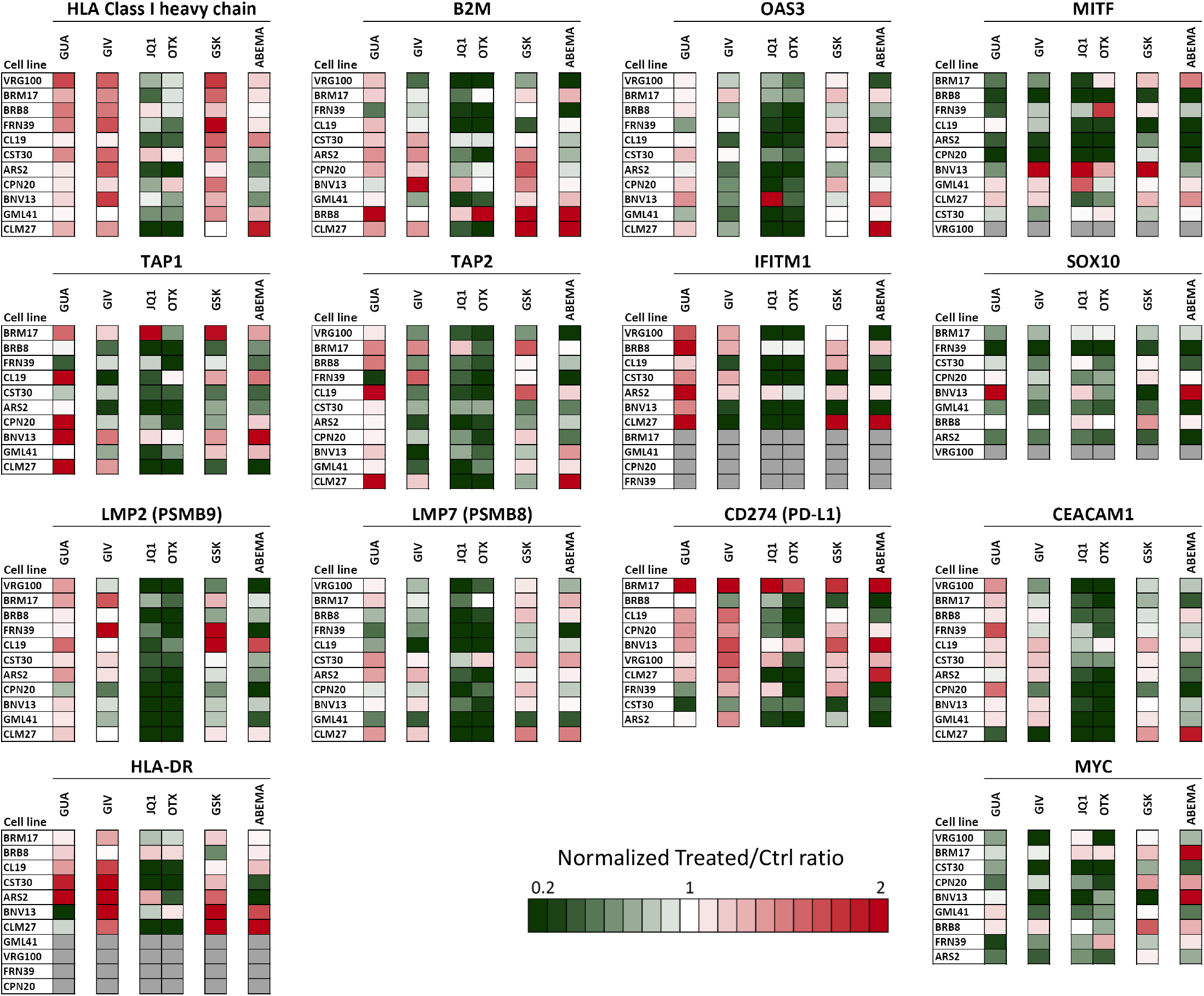
Quantitative western blot analysis for modulation of immune-related proteins in melanoma cell lines by epigenetic drugs. Eleven melanoma cell lines were treated with guadecitabine (GUA), givinostat (GIV), JQ1, OTX-015 (OTX), GSK-126 (GSK) and abemaciclib (ABEMA) and assessed for expression/modulation of 14 proteins. Modulation of each protein by drug treatment, based on data analysis and visualization approach described in Fig. S15 and S16, is visualized by a color code shown at the bottom of the figure. Grey boxes: the antigen was not expressed and not induced by treatments.

### Upstream Regulator (UR) analysis identifies activation of innate immunity and of inflammation-related pathways as the main immune-related activity of guadecitabine

To gain insight into the biological activity of the different epigenetic agents we used UR analysis, by IPA. UR analysis (pipeline of data analysis described in Fig. S17) is a computational tool that identifies the upstream transcriptional regulators that control downstream genes and explain the observed gene expression changes in the dataset. UR analysis also predicts the functional status (activated or inhibited) of each UR molecule. Each of the predicted UR can be associated with the main cellular processes and pathways it belongs, thus allowing to improve understanding of the overall biological activity of each drug (Fig. S17).

UR analysis, based on p values and activation Z scores (Table S6) and applied to gene expression dataset of cell lines VRG100 and CST30, confirmed the wide range of biological activities of all drugs. Several of the significant URs were molecules involved in immune-regulation (highlighted in green in Table S6). These UR molecules were often predicted to be activated by guadecitabine, and to a lesser extent by givinostat, but frequently inhibited by JQ1 or by GSK126. To provide a direct comparison of all drugs at the UR level, significantly modulated URs by at least two different inhibitors were grouped and visualized according to 18 functional groups (Fig. S18 and Fig. S19 dealing with cell lines VRG100 and CST30, respectively). This analysis revealed not only the breadth and drug-related specificity of epigenetic drug activity, but even the differences among these drugs (i.e. the UR with opposite activation status). A large set of these URs (group 12 in Fig. S18 and group 14 in Fig. S19) were immunity/inflammation-related molecules belonging to at least seven functional groups/pathways (cytokines, IFNs, JAK-STAT, IRFs, TLR, NF-kB, TNF) frequently activated by guadecitabine, and to a lesser extent by givinostat, while almost invariably predicted to be inhibited by JQ1. Scatter plots of significantly modulated URs by guadecitabine vs JQ1, in cell line VRG100, confirmed the opposite biological activity of these two drugs (Fig. S20A). In fact, a large set of URs activated by guadecitabine were inhibited by JQ1. In contrast, scatter plots comparing URs modulated by guadecitabine and givinostat revealed a set of URs activated by both drugs, as well as URs activated only by guadecitabine (Fig. S20B).

The UR analysis was then extended to the whole panel of cell lines using Nanostring data as input and by focusing on the activity of guadecitabine. By such analysis we identified 51 activated URs (z score >2) and 9 inhibited URs (Z score <-2) by guadecitabine in the majority of cell lines (i.e. ≥ 6/10 cell lines, Fig. 4A). Additional 36 URs were predicted to be activated in a subset of the cell lines (i.e. 5/10 cell lines, Fig. 4B). To visualize the overall biological activity of guadecitabine the URs were labeled with color-coded arrows pointing to the pathways/biological processes these molecules belong. On this basis, the main immune-related activity of guadecitabine could be summarized as an activator of innate immunity (NF-kB, TLR and Type I-III IFN) and inflammation-related pathways (Fig. 4A, B).

**Figure 4.**
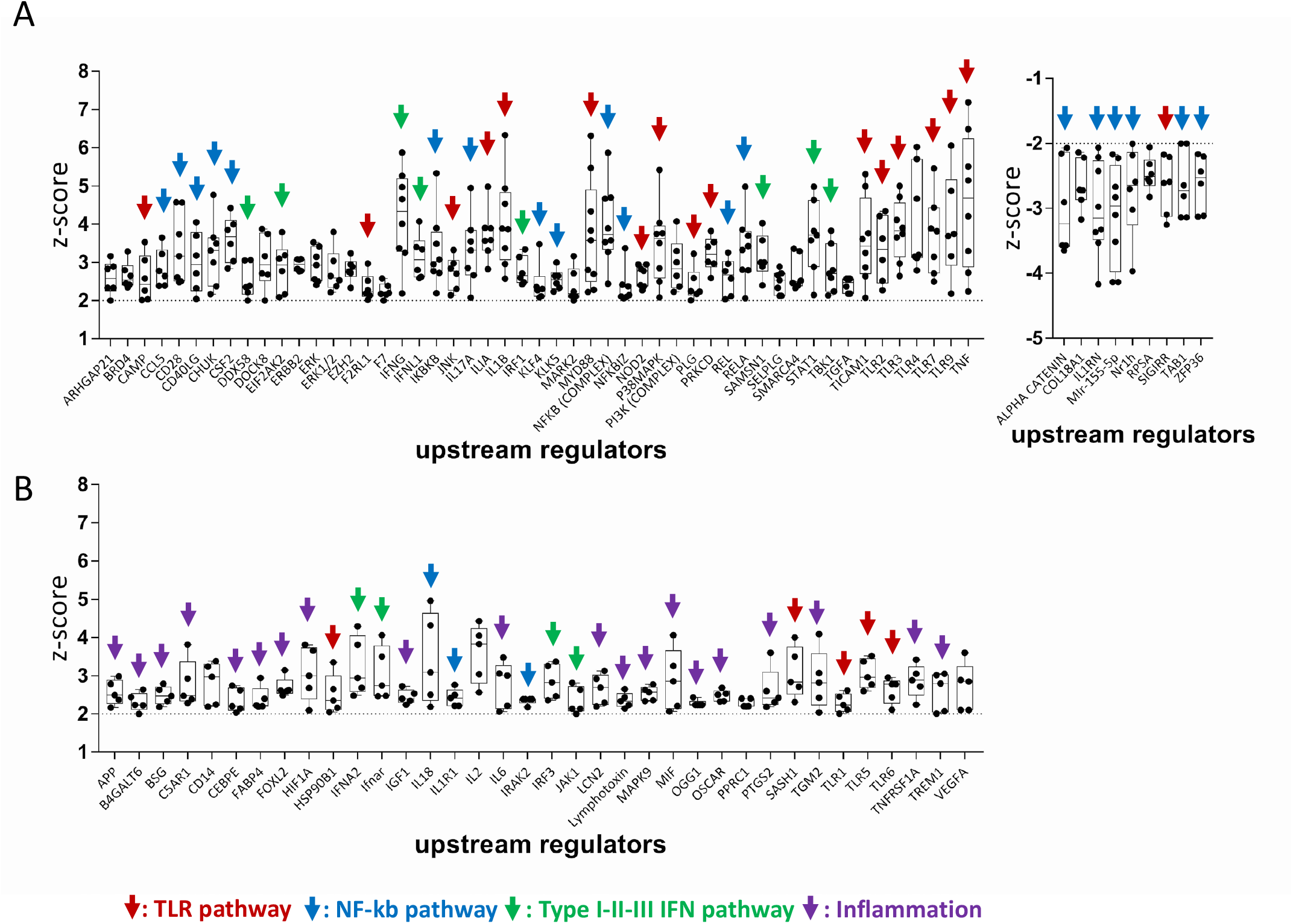
Guadecitabine is an activator of innate immunity and inflammation pathways. A. Upstream Regulators predicted to be activated (Z score >2, p value <0.05, left graph) or inhibited (Z score <-2, p value <0.05, right graph), in at least 6/10 cell lines. **B**. Upstream Regulators predicted to be activated (Z score >2, p value <0.05) in 5/10 cell lines. Data in A and B based on IPA analysis of Nanostring gene modulation data. Color-coded arrows define the biological function/pathway of each UR. Each black dot in the two graphs represents the Z score value of the indicated URs, computed for a single cell line.

We then tested whether the guadecitabine-specific UR signature could be induced by this drug in different cellular contexts, beyond the melanoma histotype, as well as in tumor nodules from a previously described [28,29] in-vivo model. By gene expression analysis in mesothelioma cell lines treated with guadecitabine and by retrieving published gene expression data of hepatocarcinoma cell lines treated with this drug [42], we found that guadecitabine-specific UR signature molecules were predicted to be activated (color-coded dots in Fig. 5A, B). Several of the guadecitabine-specific URs activated by this drug in-vitro in cell lines were also activated in-vivo in tumor nodules from immunodeficient mice bearing a human melanoma cell line xenograft [28,29] and treated with guadecitabine (Fig. S21A). In addition, canonical pathway analysis by IPA, indicated that several genes in the IFN-γ and IFN-α/β pathways were significantly upregulated by guadecitabine in the tumor xenografts (Fig. S21B). Taken together, these results indicate that guadecitabine is a strong inducer of innate immunity pathways and that its biological function can be consistently detected not only in-vitro, irrespective of tumor mutational background and histotype, but even in-vivo.

**Figure 5.**
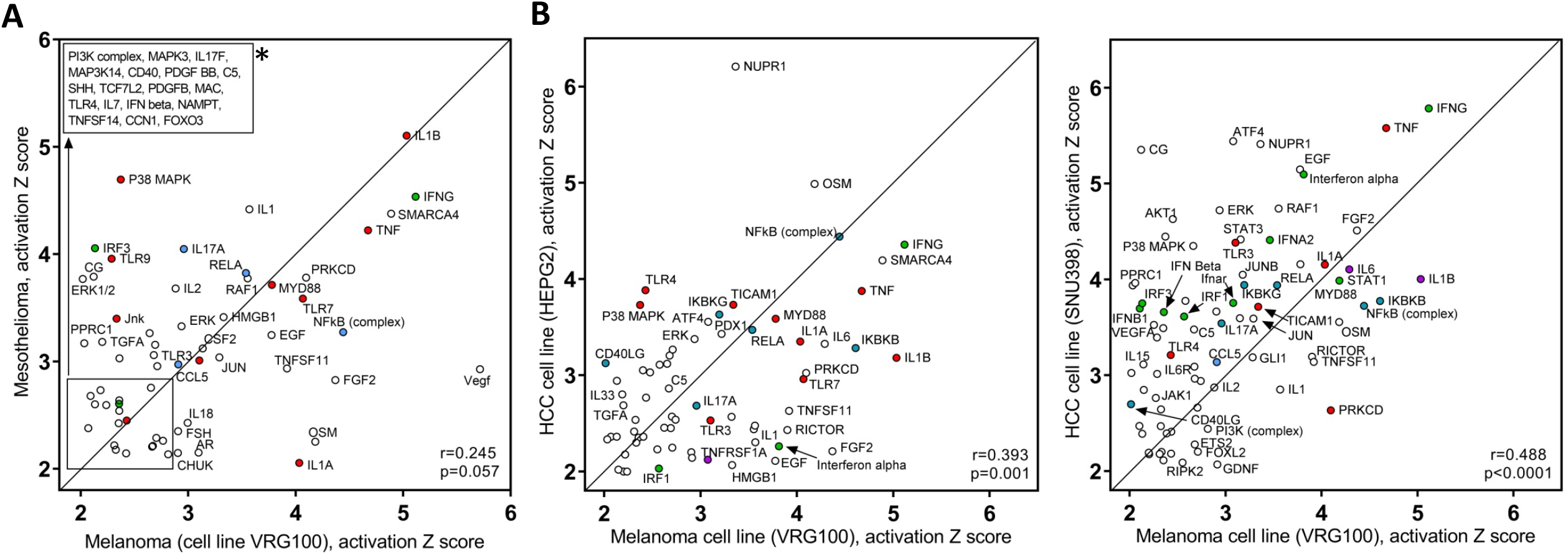
Comparison of Upstream Regulators activated by guadecitabine in melanoma vs mesothelioma and in melanoma vs hepatocarcinoma cell lines. **A, B**. Scatter plot of URs activated by guadecitabine in melanoma vs mesothelioma (A) and in melanoma vs hepatocarcinoma cell lines (B, according to gene expression data retrieved from ref. 42). Guadecitabine UR signature molecules shown in this figure are highlighted with the same color code used in Fig. 4 to mark the biological function/pathway. *: identity of URs shown in the square in panel A.

### The guadecitabine-specific gene and UR signatures are induced in-vivo in on-treatment tumor biopsies from NIBIT-M4 patients

Modulation of genes belonging to the guadecitabine-specific signature was investigated by retrieving RNA-seq data of pre-therapy (w0), and on-treatment (w4 and w12) melanoma biopsies from patients enrolled in the NIBIT-M4 trial [6]. Different groups of guadecitabine-specific signature genes (highlighted by vertical color bars in Fig. 6A) showed increased expression in w12 or w4 on-treatment samples compared to baseline or in w12 compared to w4 biopsies. Overall, almost 92% of the guadecitabine-specific signature genes were upregulated in biopsies obtained at w12 or w4, compared to w0 (Fig. 6A). By gene set enrichment analysis [43] the guadecitabine-specific gene signature showed a significant increase at w12 compared to w4 and w0, but not at w4 compared to w0 (Fig. 6B-D). We then asked whether treatment in monotherapy with the anti-CTLA-4 mAb could also lead to upregulation of most of the guadecitabine signature genes in on-treatment lesions. In two published datasets [44, 45] involving n=45 [44] and n=54 melanoma patients [45], among 376 and 286 genes upregulated by Ipilimumab in on treatment lesions compared to baseline, we found only 35/166 (21.1%) and 28/166 genes (16.9%) of the guadecitabine signature, respectively (data not shown).

**Figure 6.**
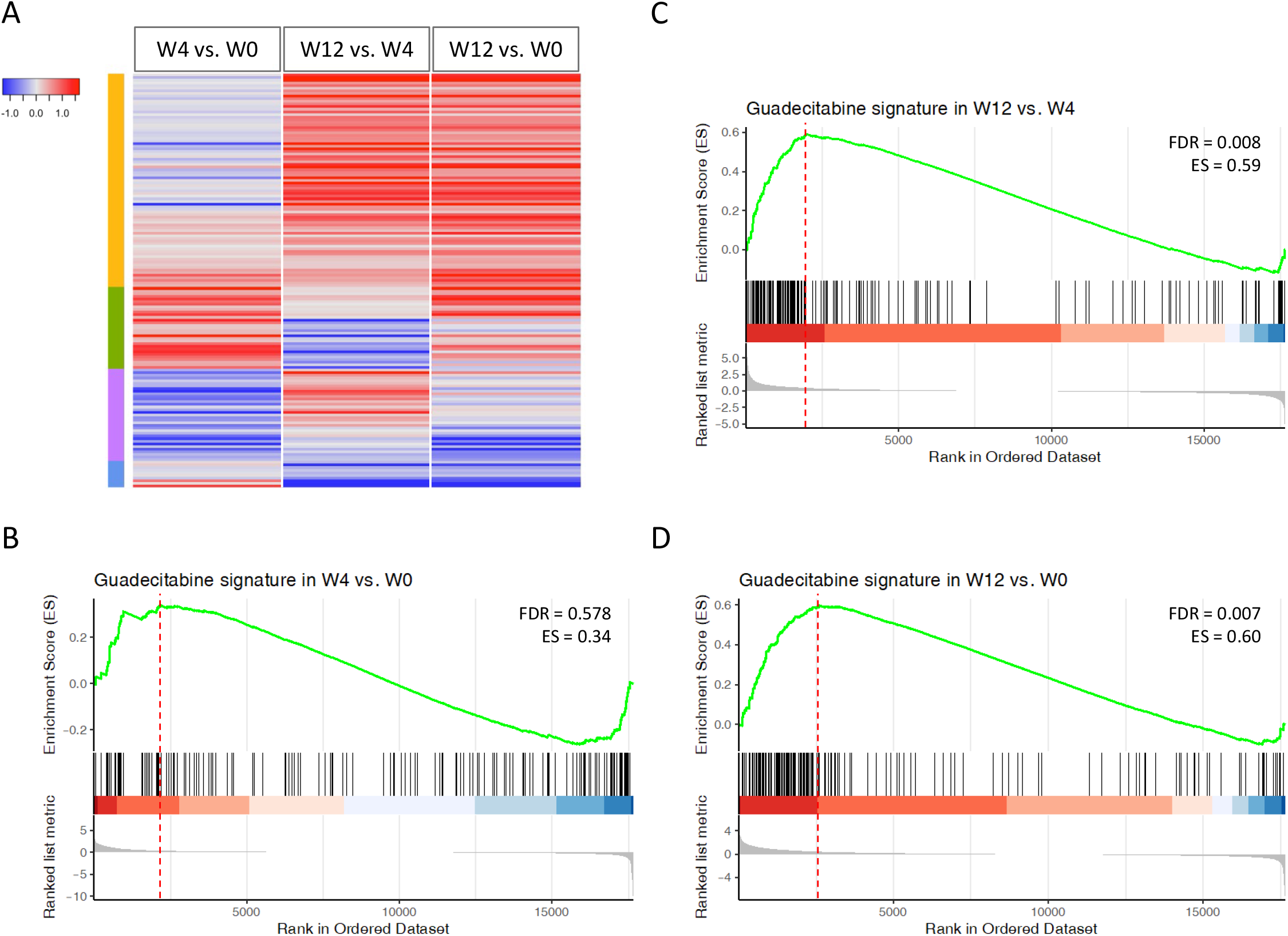
Expression and modulation of guadecitabine-specific signature genes in baseline and on-treatment clinical samples from NIBIT-M4 patients. **A**. Heatmap of log2 fold changes of guadecitabine signature genes among the following comparisons: w4 vs. w0, w12 vs. w4 and w12 vs w0. **B-D**. Enrichment plots containing profiles of the running enrichment scores (ES) and positions of guadecitabine gene set members on the rank ordered list in GSEA for w4 vs. w0 (B), w12 vs. w4 (C) and w12 vs. w0 (D) comparisons.

We then identified genes showing differential expression, at each time point, in biopsies from responding vs non-responding patients of the NIBIT–M4 trial and then we subjected these differentially expressed genes to UR analysis by IPA. The results revealed progressive activation of several guadecitabine-specific UR signature molecules in on-treatment (w4 and w12) tumor biopsies from responding compared to non-responding patients (Fig. 7A, B), as well as a strong correlation of these activated UR at w4 vs w12 tumor samples (Fig. 7C).

**Figure 7.**
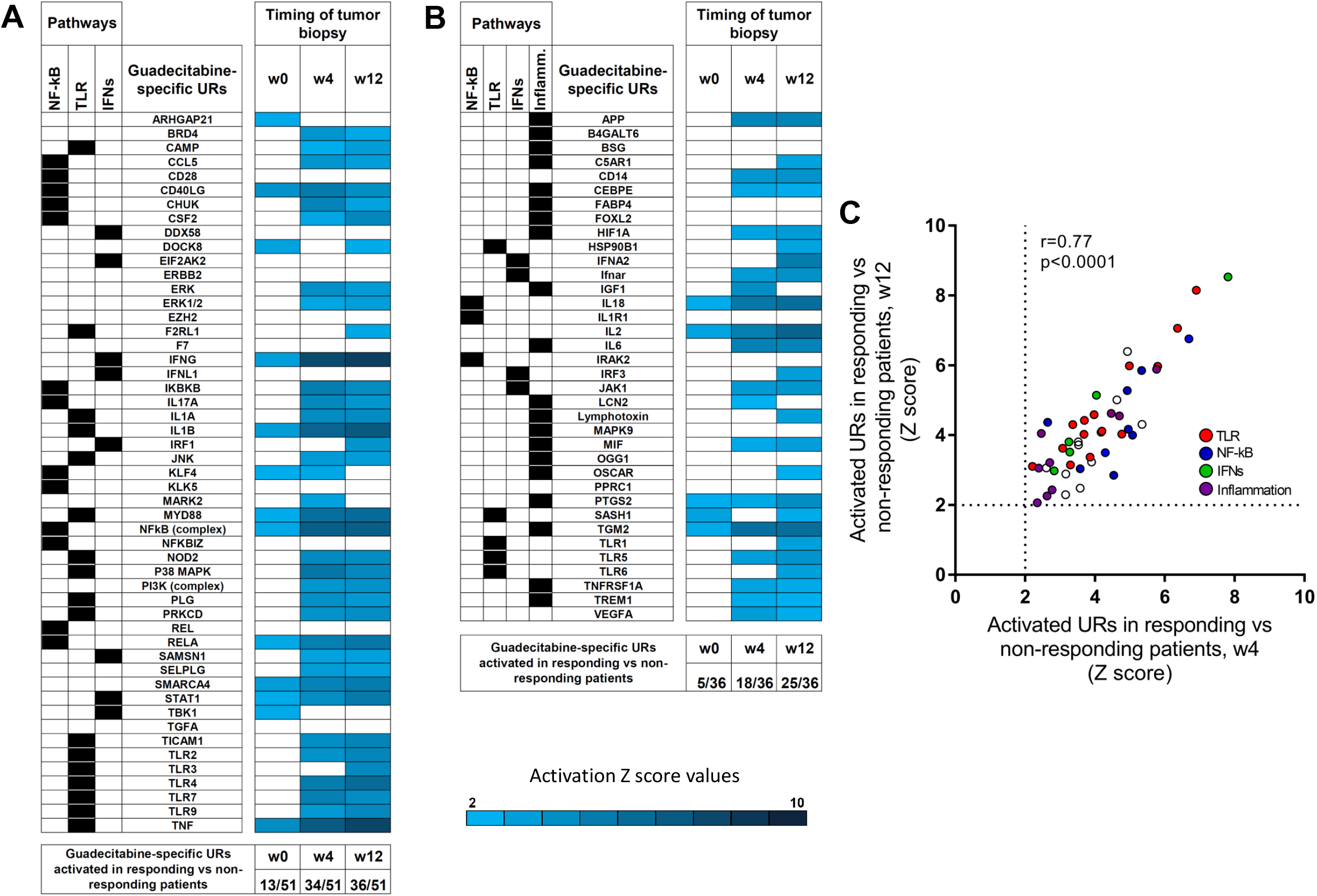
Activation of guadecitabine-specific UR signature molecules in on-treatment tumor biopsies from responding compared to non-responding patients in the NIBIT-M4 trial. **A**, B. Z score values of guadecitabine-activated URs as identified in Fig. 4A (A) or in Fig. 4B (B) based on differential gene expression analysis in tumor samples (obtained at w0, w4 and w12) from responding (n=3) vs non-responding patients (n=5). Only significant URs (Z score >2) are shown by the indicated color code. **C**. Scatter plot of Z score values of activated URs in responding vs non-responding patients at w4 vs. wk12.

Taken together these results strongly support the notion that the guadecitabine-specific gene signature defined in-vitro in melanoma cell lines can be modulated even in-vivo at tumor site in patients treated with this DNMT inhibitor. Moreover, the progressive activation of the UR signature in responding vs non responding patients suggests that the clinical activity of guadecitabine in melanoma patients depends on effective activation of innate immunity pathways.

## Discussion

Characterization of the biological activity of the four investigated agents targeting distinct epigenetic regulators by gene expression, quantitative western blot and UR analyses in melanoma cell lines, indicated that the DNMT inhibitor guadecitabine and the BET inhibitor JQ1 were the most active drugs, but exerted opposite immuno-regulatory functions. Comparison of the drug-specific immune-related gene signatures indicated that guadecitabine upregulated several genes in each of the 21 functional classes present on the Nanostring Cancer Immune panel, while JQ1 predominantly downregulated genes in such classes. Moreover, almost all immune-related URs predicted to be activated by guadecitabine were instead inhibited by JQ1 in the majority of the melanoma cell lines tested. Among UR predicted to be inhibited by JQ1 we found not only molecules belonging to the IFN-gamma pathway, known to be targeted by this BET inhibitor [46], but also molecules belonging to the TLR and NF-kb pathways regulated by the BET proteins [21]. The experiments looking at gene modulation in melanoma cell lines, by treatment with combinations of two different epigenetic inhibitors, confirmed that the guadecitabine and JQ1 were the dominant drugs that determined opposite responses to treatment, irrespective of the other drug used in the combination. Collectively, the results suggest that guadecitabine elicits marked pro-inflammatory effects, while JQ1 is mainly an anti-inflammatory drug.

The breadth of non-immuno-related effects of the four epigenetic inhibitors emerged through comparative transcriptomic analysis. The four drugs had a variable impact on expression of genes encoding DNMT, HDAC, histones, BET, and PRC2 components. This suggests that drug activity of these agents is not uniquely dependent on inhibition of their specific targets, but even on modulation of additional genes which contribute to different aspects of epigenetic regulation. The drugs affected also expression of genes encoding proteolytic enzymes (cathepsins), and their inhibitors (serpins), thus implicating epigenetic drugs in the regulation of different cancer-related processes, controlled by cathepsins, such as proliferation, angiogenesis, metastasis and invasion. Taken together, these results significantly extend previous knowledge on non-immuno-related anti-tumor activities of epigenetic inhibitors, such as reactivation of tumor suppressor genes by demethylating agents [47], promotion of cell death and suppression of angiogenesis by HDAC inhibitors [48], inhibition of proto-oncogenes MYC and BCL2 by BET inhibitors [49] and downregulation of DNA repair genes by EZH2 inhibitors [50].

In depth analysis of guadecitabine biological activity indicated that this DNMT inhibitor upregulated genes belonging to 21 immune-related classes. The >160-gene drug-specific signature of guadecitabine and the related UR signature profiled the remarkable immunomodulatory activity of this drug. The emerging immune-related biological activity of guadecitabine, identified through UR analysis, was consistently observed across a panel of melanoma cells lines with widely divergent mutational and differentiation profiles, as well as across different tumor histotypes. A pre-clinical tumor model confirmed the activation of the guadecitabine-specific UR signature even in-vivo. Finally, we found that most of the guadecitabine-specific signature genes were upregulated at w4 or w12 in clinical samples from NIBIT-4 patients, but not in on treatment tumor samples from patients treated with Ipilimumab only [44-45]. Moreover, progressive activation of the guadecitabine-specific UR signature could distinguish on treatment tumor biopsies of responding compared to non-responding patients. These results suggest that the clinical efficacy of guadecitabine in melanoma patients is strictly associated with the effective promotion, in-vivo, of the innate immunity pathways that define the biological activity of this DNMT inhibitor.

In conclusions, the findings of this study support a mechanistic rationale for development of combinatorial immune intervention where boosting of NF-kB, TLR, Type I-III IFN and inflammation-related pathways by the DNMT inhibitor guadecitabine may cooperate with rescue of adaptive immunity by immune checkpoint blockade to improve efficacy of cancer immunotherapy.

## Supporting information

Supplementary Sable S1

## Abbreviations

APM: antigen processing machinery
BET: bromodomain and extra-terminal domain
CT: cancer testis
DNMT: DNA methyltransferases
EZH2: enhancer of zeste homolog 2
HDAC: histone deacetylases
ICI: immune checkpoint inhibitor
IPA: Ingenuity Pathway Analysis
LSD-1: Lysine-specific demethylase 1
MTT: 3-(4,5)dimethylthiazol-2,5-diphenyltetrazolium bromide
NIBIT: Italian Network for Tumor Biotherapy
NGS: next generation sequencing
TAC: Transcriptomic Analysis Console
UR: Upstream Regulator.

## Supplementary Information

**-Supplementary Methods and Figures.pdf**. This file contains supplementary materials and methods and supplementary figures S1 to S21. **-Fig. S1**. Expression in ten melanoma cell lines of genes and gene families targeted by epigenetic drugs. **-Fig. S2**. Melanoma differentiation profile of cell lines according to expression of seven subtype signatures and four main melanoma subsets. **-Fig. S3**. Susceptibility of ten melanoma cell lines to the anti-proliferative effects of epigenetic drugs. **-Fig. S4**. Volcano plot of differentially expressed genes in VRG100 and CST30 cell lines. **-Fig. S5**. Whole genome gene modulation analysis by epigenetic drugs in two melanoma cell lines. -**Fig. S6**. Edwards-VENN diagram analysis of significantly modulated genes. **-Fig. S7**. Original and revised gene classification of the NanoString nCounter PanCancer Immune Profiling panel. **-Fig. S8**. Quantitative analysis of Nanostring data in ten melanoma cell lines treated with epigenetic drugs. **-Fig. S9**. Modulation of immune-related genes in melanoma cell lines by epigenetic drugs. **-Fig. S10**. Comparison of immune-related gene modulation by BET inhibitors JQ1 and OTX-015. **- Fig. S11**. Immune-related signatures of epigenetic drugs in melanoma. **-Fig. S12**. Expression/methylation relationship, for selected genes in the guadecitabine-specific gene signature, among melanoma cell lines present in the GDSC-MGH Sanger database. **-Fig. S13**. Expression/methylation relationship of genes belonging to the guadecitabine-specific signature according to the GDSC-MGH Sanger melanoma dataset. **-Fig. S14**. Changes in gene expression and gene methylation for the cancer testis class of genes in tumor biopsies from melanoma patients enrolled in the NIBIT-M4 trial. **-Fig. S15**. Outline of the strategy for quantitative analysis and visualization of western blot data. **-Fig. S16**. Quantitative western blot analysis and visualization of the modulation of LMP7 by epigenetic drugs in 11 melanoma cell lines. **-Fig. S17**. Pipeline of data analysis based on Upstream Regulators (UR) identified by IPA. **-Fig. S18**. Classification of URs significantly modulated by at least two different drugs in melanoma cell line VRG100. **-Fig. S19**. Classification of URs significantly modulated by at least two different drugs in melanoma cell line CST30. **-Fig. S20**. Comparison of all URs predicted to be activated or inhibited by Guadecitabine vs JQ1 and by Guadecitabine vs Givinostat. **-Fig. S21**. Comparison of URs activated by guadecitabine in-vitro and in-vivo.

**-Table S1.xlsx**. Origin of melanoma cell lines, histopathological features of the corresponding primary tumor, cell line authentication data by STR profiling, and mutational profile by NGS of 14 melanoma cell lines used in this study.

**-Table S2.xlsx**. Drug doses used for experiments and effect of each drug at such dose in the MTT assay on each cell line.

**-Table S3.xlsx**. Antibodies used in quantitative western blot analysis.

**-Table S4.xlsx**. Modulation of immune-related genes belonging to 21 functional classes by epigenetic drugs. This excel file has 21 sheets.

**-Table S5.xlsx**. Immune-related gene modulation by epigenetic drugs and by combinatorial treatments. This excel file has 5 sheets.

**-Table S6.xlsx**. Upstream Regulator analysis on differentially expressed genes in drug-treated vs untreated cell lines VRG100 and CST30.

## Declarations

### Ethics approval and consent to participate

The data analyzed during this study concern gene expression and methylation in tumor samples from the published NIBIT-M4 trial [6]. The NIBIT-M4 Phase Ib trial was conducted in accordance with the ethical principles of the Declaration of Helsinki and the International Conference on Harmonization of Good Clinical Practice. The protocol was approved by the independent ethics committee of the University Hospital of Siena (Siena, Italy). All participating patients (or their legal representatives) provided signed-informed consent before enrolment. The trial was registered with European Union Drug Regulating Authorities ClinicalTrials EudraCT, number 2015-001329-17, and with ClinicalTrials.gov, number NCT02608437.

### Consent for publication

Not applicable.

### Availability of data and materials

The datasets generated during the current study are available in the Gene Expression Omnibus (GEO) repository with the following GEO accession numbers: GSE189628, GSE189629, GSE189630, GSE189631. The data analyzed during this study concerning gene expression and methylation in tumor samples from patients enrolled in the published NIBIT- M4 trial were retrieved from ref. 6.

### Competing interests

AA has received compensated educational activities from Bristol-Myers Squibb. AMDG has served as a consultant and/or advisor to Incyte, Pierre Fabre, Glaxo Smith Kline, Bristol-Myers Squibb, Merck Sharp Dohme, and Sanofi and has received compensated educational activities from Bristol Myers Squibb, Merck Sharp Dohme, Pierre Fabre and Sanofi. MM has served as a consultant and/or advisor to Roche, Bristol-Myers Squibb, Merck Sharp Dohme, Incyte, AstraZeneca, Amgen, Pierre Fabre, Eli Lilly, Glaxo Smith Kline, Sciclone, Sanofi, Alfasigma, and Merck Serono; and owns shares in Epigen Therapeutics. SC and AC own shares in Epigen Therapeutics. G.Palmieri has advisory role for Bristol-Myers Squibb, Merck Sharp Dohme, Roche, Novartis, Pierre-Fabre, Incyte. A.Maurichi has received compensated educational activities from Novartis. G.Pruneri has received meetings honoraria from from AstraZeneca, Novartis, Exact Sciences, Roche and Illumina and is advisory board member of ADS Biotec. A.Molla, GN, VEP, FS, CF, MFL, A.Manca, MCS, MP, SB, TN,FC, MS and RM declare no conflicts of interests.

## Funding

The research leading to these results has received funding from: AIRC under 5 per mille 2018 – ID.21073 program – P.I. Maio Michele, Group Leader Anichini Andrea and Ministry of Health, Lombardy and Tuscany regions, Bando Ricerca Finalizzata, grant number NET-2016- 02361632, P.I. Michele Maio.

## Authors’ contributions AA

conceptualization, funding acquisition, resources, supervision, data curation, formal analysis, visualization, methodology, writing of original draft. A.Molla: cell biology experiments, methodology, data curation. GN: molecular biology experiments, methodology, data curation. VEP: western blot experiments, methodology, formal analysis, data curation. FS: cell biology experiments, methodology, visualization, data curation. AC: data curation, methodology, writing–review and editing. CF: in-vivo and in-vitro experiments, methodology, data curation. MFL: in-vivo and in-vitro experiments, methodology, data curation. AMDG: data curation, resources, writing–review and editing. SC: Project administration, data curation, resources. A.Manca: NGS experiments, methodology, data analysis and curation. MCS: NGS experiments, data analysis and curation, methodology. MP: NGS experiments, methodology, data analysis and curation. TN: data analysis; FC: data analysis. SB: Nanostring experiments, methodology, data curation. G.Pruneri: resources, writing–review and editing. A.Maurichi: data curation, writing–review and editing. MS: data curation, writing–review and editing. MC: conceptualization, data analysis, writing–review and editing. G.Palmieri: conceptualization, funding acquisition, data curation, resources, formal analysis. MM: conceptualization, funding acquisition, data curation, resources, writing–review and editing. RM: conceptualization, supervision, data analysis and curation, writing of original draft.

## Acknowledgements

The authors wish to tank the **EP**igenetic **I**mmune-oncology **C**onsortium **A**IRC (**EPICA**) investigators (Daniela Massi, Ulrich Pfeffer) for helpful discussion of data and critical review of the manuscript.

## Notes

### Competing Interest Statement

A.Anichini has received compensated educational activities from Bristol-Myers Squibb. A.M.Di Giacomo has served as a consultant and/or advisor to Incyte, Pierre Fabre, Glaxo Smith Kline, Bristol-Myers Squibb, Merck Sharp Dohme, and Sanofi and has received compensated educational activities from Bristol Myers Squibb, Merck Sharp Dohme, Pierre Fabre and Sanofi. M.Maio has served as a consultant and/or advisor to Roche, Bristol-Myers Squibb, Merck Sharp Dohme, Incyte, AstraZeneca, Amgen, Pierre Fabre, Eli Lilly, Glaxo Smith Kline, Sciclone, Sanofi, Alfasigma, and Merck Serono; and owns shares in Epigen Therapeutics. S.Coral and A.Covre own shares in Epigen Therapeutics. G.Palmieri has advisory role for Bristol-Myers Squibb, Merck Sharp Dohme, Roche, Novartis, Pierre-Fabre, Incyte. A.Maurichi has received compensated educational activities from Novartis. G.Pruneri has received meetings honoraria from from AstraZeneca, Novartis, Exact Sciences, Roche and Illumina and is advisory board member of ADS Biotec. A.Molla, G.Nicolini, V.E.Perotti, F.Sgambelluri, C.Fazio, M.F.Lofiego, A.Manca, M.C.Sini, M.Pisano, S.Brich, T.Noviello, F.Caruso, M.Santinami and R.Mortarini declare no conflicts of interests.

