## Supplementary Methods and Supplementary Figures for "Landscape of immune-related signatures induced by targeting of different epigenetic regulators in melanoma: implications for immunotherapy"

On behalf of the **EP**igenetic **I**mmune-oncology **C**onsortium **A**IRC (**EPICA**) investigators.

[1]Human Tumors Immunobiology Unit, Dept. of Research, [7]Department of Pathology and Laboratory Medicine, [9]Melanoma and Sarcoma Unit, Department of Surgery, Fondazione IRCCS Istituto Nazionale dei Tumori, Milan, Italy. [2]Center for Immuno-Oncology, University Hospital of Siena, Siena, Italy. [3]University of Siena, Siena, Italy. [4]Unit of Cancer Genetics, National Research Council (CNR), Sassari, Italy. [10]University of Sassari, Sassari, Italy. [5]Department of Electrical Engineering and Information Technology (DIETI), University of Naples "Federico II", Naples, Italy. [6]BIOGEM Institute of Molecular Biology and Genetics, Ariano Irpino, Italy. [8]University of Milan, School of Medicine, Italy.
*These authors contributed equally.

**Correspondence**: Andrea Anichini, PhD, Human Tumors Immunobiology Unit, Dept. of Research, Fondazione IRCCS Istituto Nazionale dei Tumori, Via Venezian 1, 20133 Milan, Italy. Phone+390223902817..

#### Supplementary Methods.

**NGS analysis.** Next generation sequencing (NGS) assays on melanoma cell lines DNA were performed using Ion GeneStudio S5 System and carried out on Ion AmpliSeq™ Comprehensive Cancer Panel, which provides highly multiplexed target selection of 409 genes implicated in cancer pathogenesis. Starting DNA and libraries were accurately quantified using a fluorescence-based method, such as Qubit dsDNA HS. Data analysis workflow was performed by automated data transfer, from the Ion Torrent™ Server to the Ion Reporter Server for variant analysis; it includes result filtering, annotation, and data analysis results. To get a total amount of at least 10 mutated alleles for each candidate amplicon, the following mutation selection criteria were adopted:

coverage of >200 reads and frequency of mutated alleles >5% for gene amplicon. The copy number variation (CNV) determination was obtained by adding a custom control copy number baseline to the comprehensive cancer profile analysis workflow. Results of NGS analysis of 14 melanoma cell lines used in this study are shown in Supplementary Table S1D**.**

**Gene expression analysis.** The total RNA (20 ng to 50 ng) was reverse transcribed using GeneChip® WT Pico Reagent Kit (Affymetrix; Thermo Fisher Scientific, Inc.). The resulting cDNA was used as a template for in vitro transcription using the same kit. The obtained antisense cRNA was purified using Nucleic Acid Binding Beads (GeneChip® WT Pico Reagent Kit, Affymetrix) and used as a template for reverse transcription to produce single-stranded DNA in the sense orientation. During this step, dUTP was incorporated. The DNA was then fragmented using uracyl DNA glycosylase (UDG) and apurinic/apyrimidinic endonuclease 1 (APE 1) and labeled with DNA reagent covalently linked to biotin using terminal deoxynucleotidyl transferase (TdT, GeneChip® WT Pico Reagent Kit, Affymetrix). Hybridization of each fragmented and labeled target was performed using the GeneChip® Hybridization, Wash and Stain Kit (Affymetrix; Thermo Fisher Scientific, Inc). A single GeneChip® Human Clariom S was then hybridized with each biotin-labeled sense target. GeneChip arrays were scanned using an Affymetrix GeneChip® Scanner 3000 7G using default parameters. Affymetrix GeneChip® Command Console software (AGCC) was used to acquire GeneChip® images and generate .DAT and .CEL files. Gene expression data were analyzed by Transcriptomic Analysis Console (TAC) software (Applied Biosystems, Thermo Fisher Scientific). Modulation of immune-related genes by epigenetic drugs in ten melanoma cells was assessed by the NanoString nCounter PanCancer Immune profiling panel enabling determination of 731 genes (NanoString Technologies, Seattle, USA). The manufacturer's gene classification associated with the PanCancer Immune profiling panel was revised by retrieving information on gene function at http://genecards.org and then by grouping genes into 21 functional classes. For Nanostring experiments panel probes (capture and report) and 200 ng of RNA were hybridized overnight at 65 °C for 16 h. Samples were scanned at maximum scan resolution capabilities (555 FOV) using the nCounter Digital Analyzer. Quality control of samples, data normalization and data analysis were performed using nSolver software 4.0 (NanoString Technologies).

Whole gene expression profile of treated and untreated mesothelioma cell lines was performed by Agilent whole human genome oligo microarray kits. The quantity and the quality of RNA, extracted as previous described, was assessed with NanoDrop® ND-1000 UV-Vis Spectrophotometer (NanoDrop Technologies, Wilmington, DE, USA) and the Agilent 2100 Bioanalyzer (Agilent Technologies, Santa Clara, CA, USA). *In vitro* transcription, labeling and purification of dye-labeled cRNA were performed using the Quick Amp Labeling Kit, one-color (Agilent Technologies, Santa Clara, CA, USA) following manufacturer's guidelines. Gene expression profiling was performed by a One-Color strategy using Cy3-labeled aRNA from guadecitabine-treated and untreated cells (Quick Amp Labeling, Agilent Technologies, Santa Clara, CA, USA). A mixture of 1650 ng of Cy3-labeled reference cRNA, Blocking Agent and Fragmentation Agent was hybridized to Whole Human Genome (1x44K) oligo microarray platform (Agilent Technologies, Santa Clara, CA, USA). Hybridization was performed for 17 hours at 65°C in 2x GEx Hybridation Buffer HI-RPM (Agilent Technologies, Santa Clara, CA, USA), using Agilent's Hybridization Oven at 10 rpm. Following washing, slides were analyzed by Agilent Microarray Scanner. Feature Extraction Software provided by Agilent (version 9.5.3) was used to quantify the intensity of fluorescence images and to normalize results by subtracting local background fluorescence according to the manufacturer's instruction. Genes modulated with a FC ≥ 2 or ≤ -2 in treated vs untreated cells were used for upstream regulator analyses.

**Ingenuity Pathway Analysis (IPA).** Upstream regulator analysis allows to identify upstream transcriptional regulators that can explain the observed gene expression changes in the dataset.

This computational tool returns results based on p-values and Z score statistics. P values indicate the likelihood of the association between a set of genes and related function, or the likelihood of the overlap between the genes in the dataset and those that are regulated by a predicted upstream regulator. The meaning of the Z score statistics is to infer the activation states (“increased” or “decreased”) of the identified biological functions and of the predicted transcription factors. Only Z scores greater than 2 or smaller than -2 were considered significant. Canonical pathway analysis is a computational tool allowing to determine if canonical pathways are activated or inhibited on the basis of gene expression in the dataset. Activation or inhibition states of canonical pathways are predicted based on the Z-score algorithm. The significance values (p-value of overlap) are calculated by the right-tailed Fisher's Exact Test and indicate the probability of association of molecules in the dataset with the canonical pathway by random chance alone.

**Relationship of gene expression with promoter methylation**. Data on gene expression and promoter methylation ($\beta$ values) of melanoma cancer cell lines in the GDSC-MGH Sanger database were retrieved from CellMiner CDB web site (at: https://discover.nci.nih.gov/cellminercdb/). Clustering of gene expression data was carried out by Cluster 3.0.

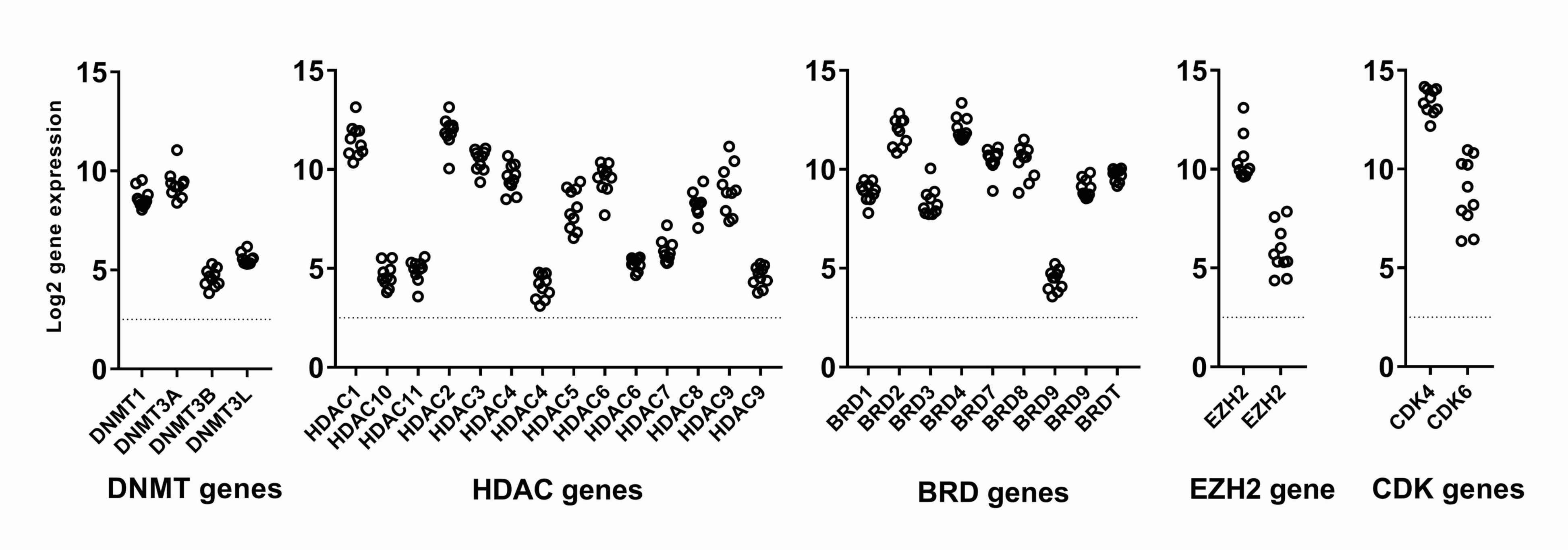


**Supplementary Figure S1**. Expression in ten melanoma cell lines of genes and gene families targeted by Decitabine/Guadecitabine (DNMT genes), Givinostat (HDAC genes), JQ1 or OTX-015 (BRD genes), GSK-126 (EZH2 gene, two different probes present for this gene in the Clariom S array), and Abemaciclib (CDK4/6 genes). Melanoma cells lines tested in this assay were: GML41, VRG100, BRM17, CST30, BNV13, CPN20, FRN39, GRD43, SLN91, MNT59.

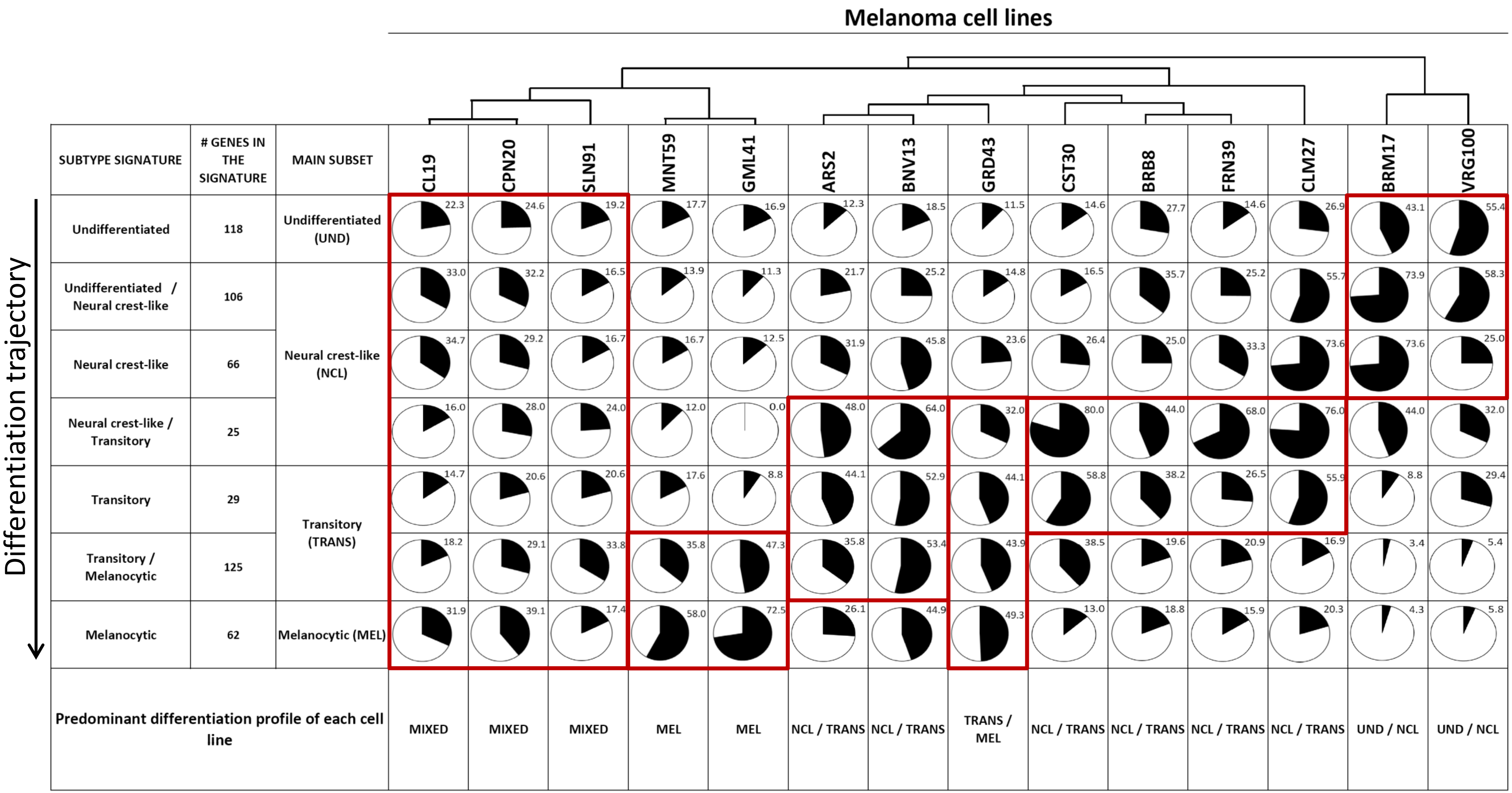


**Supplementary Figure S2. Melanoma differentiation profile of cell lines according to expression of seven subtype signatures and four main melanoma subsets as defined by Tsoi et al (36).** Expression of all genes belonging to each of seven subtype signatures was evaluated by Clariom S arrays. For each cell line the pie charts indicate the % of genes within each subtype signature that have median centered expression >0.5 (in Log2 space).
Red rectangles highlight the predominant differentiation profile of each cell line. MEL: melanocytic; NCL/TRANS: neural crest-like / transitory; TRANS/MEL: transitory / melanocytic; UND/NCL: undifferentiated / neural-crest-like. Cell lines were clustered by Cluster 3.0 according to the expression of genes in each subtype signature.

Supplementary Figure S3

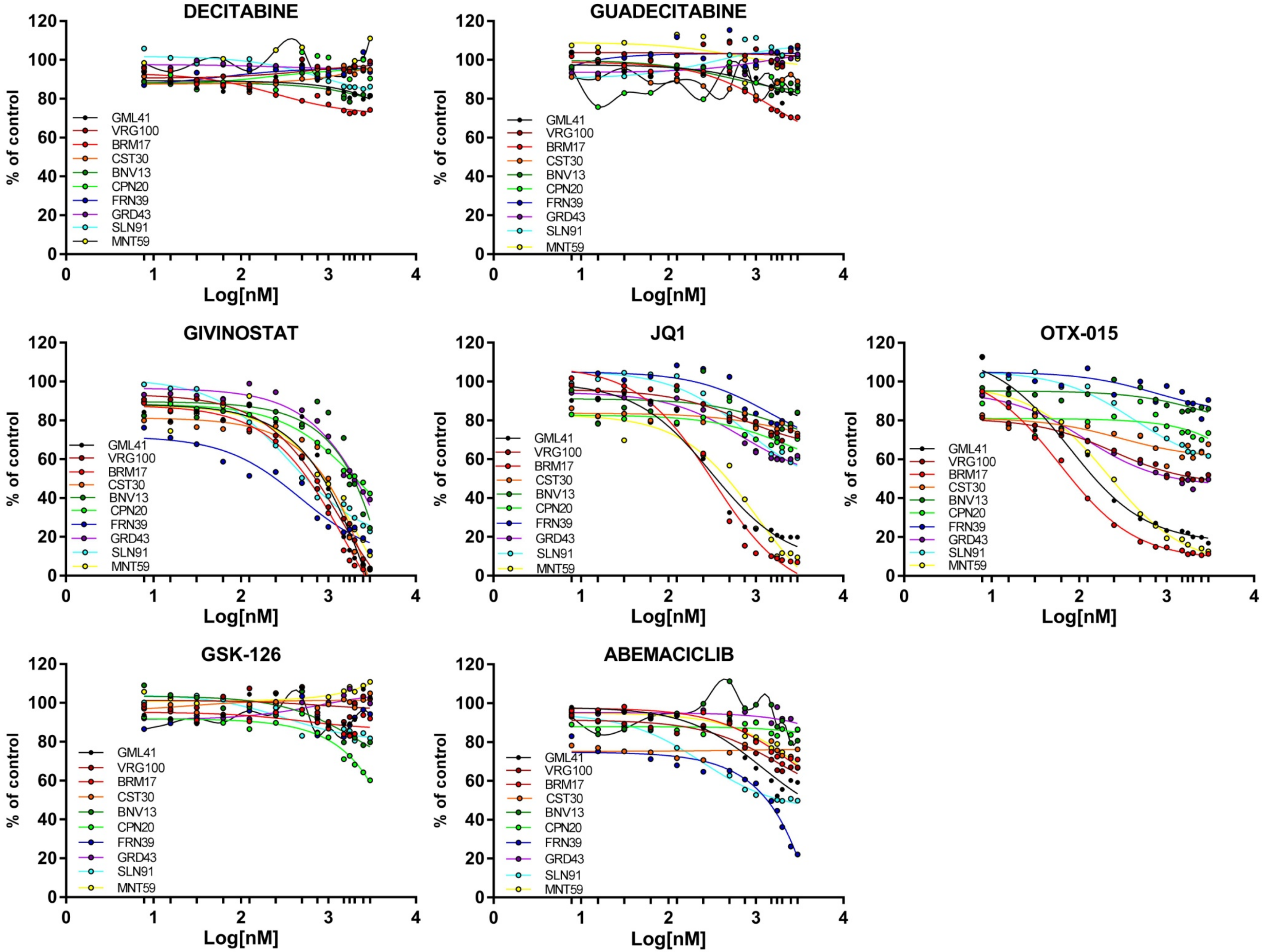


**Supplementary Figure S3**. Susceptibility of ten melanoma cell lines to the anti-proliferative effects of the indicated epigenetic drugs was evaluated at 96h by the MTT assays.

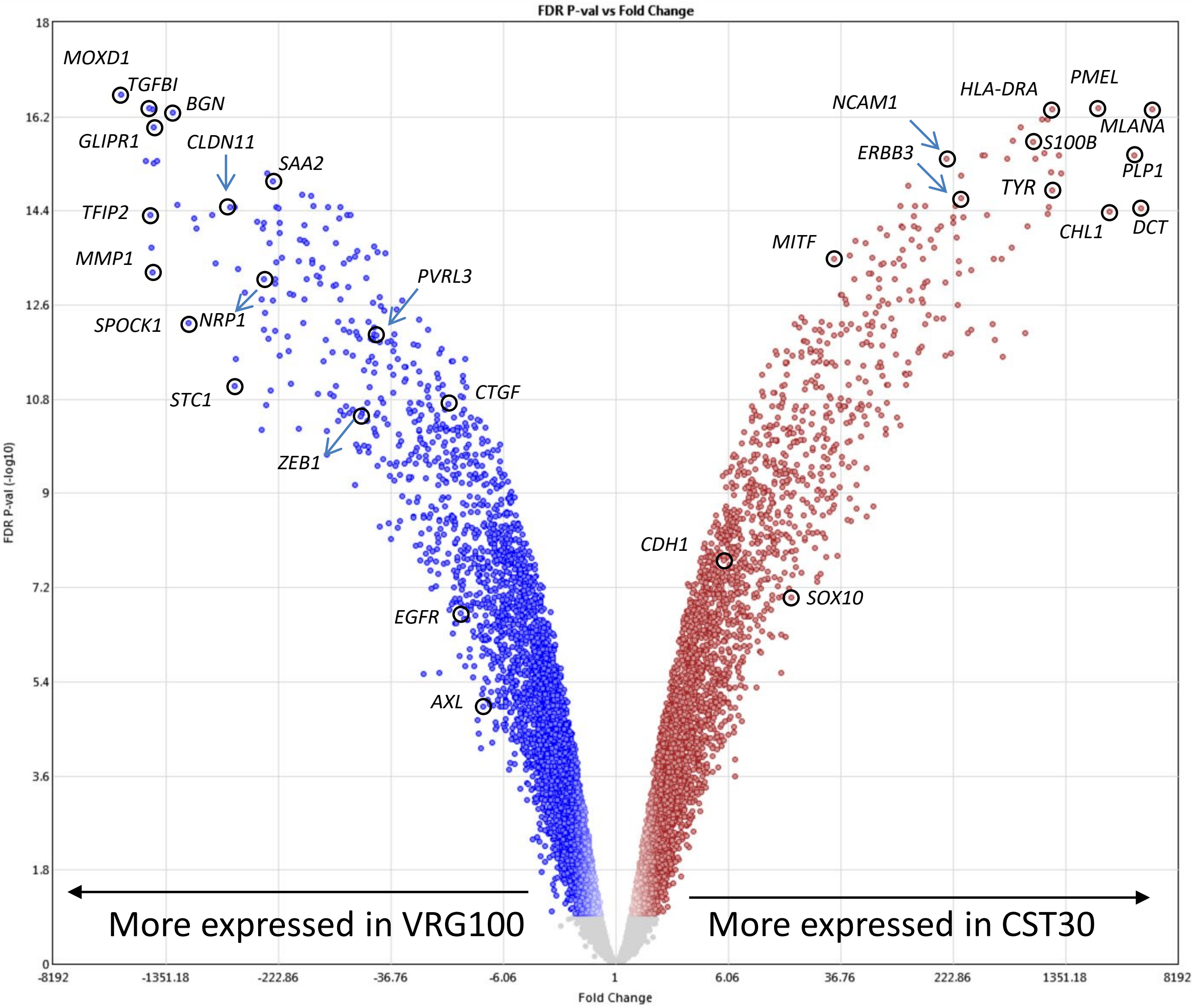


**Supplementary Figure S4**. Volcano plot of differentially expressed genes in VRG100 and CST30 cell lines. Genes identified by circles and gene symbols highlight the divergent phenotypic profile of the two cell lines, with CST30 showing higher expression of several genes associated with a more differentiated state (e.g. *MITF, SOX10, PMEL, MLANA, TYR, DCT, ERBB3*) and VRG100 showing higher expression of genes associated with a more undifferentiated/mesenchymal state (e.g. *AXL, EGFR, ZEB1, TGFBI, SPOCK1, PVRL3, CTGF ).*

Supplementary Figure S5

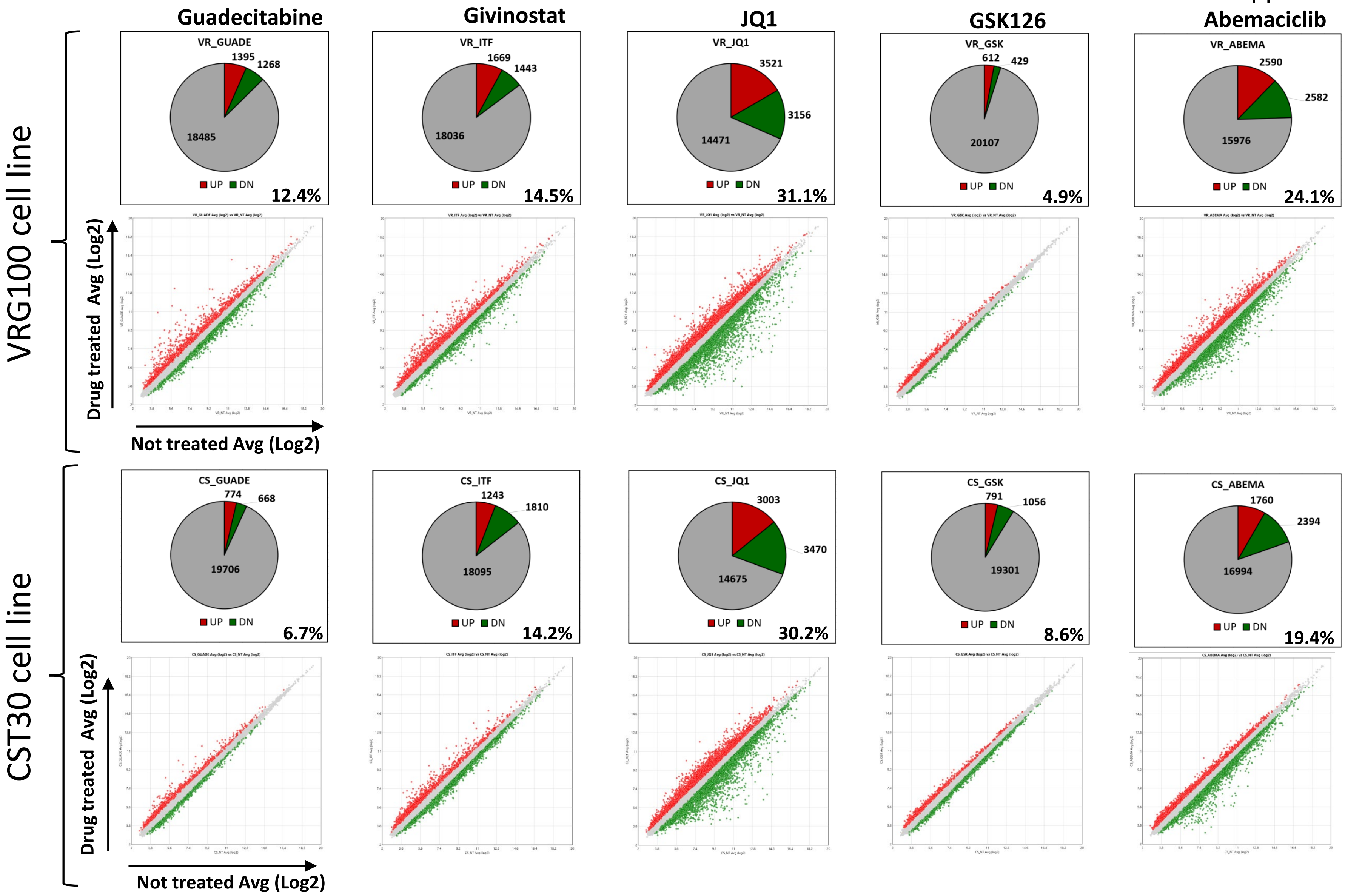


**Supplementary Figure S5**. Whole genome gene modulation analysis by 4 epigenetic drugs (guadecitabine, givinostat, JQ1, GSK126) and by a control drug (abemaciclib) in two melanoma cell lines (top graphs: VRG100; bottom graphs: CST30). For each cell line and drug, quantitative data of gene modulation are shown as pie charts indicating the number of up-regulated (red), down-regulated (green) or not modulated (grey) genes and the % of genes passing the filter (FC |>1.2|, p<0.05). Scatter plots show extent of gene modulation by each drug (red: upregulated genes, green: downmodulated genes) in the two cell lines.

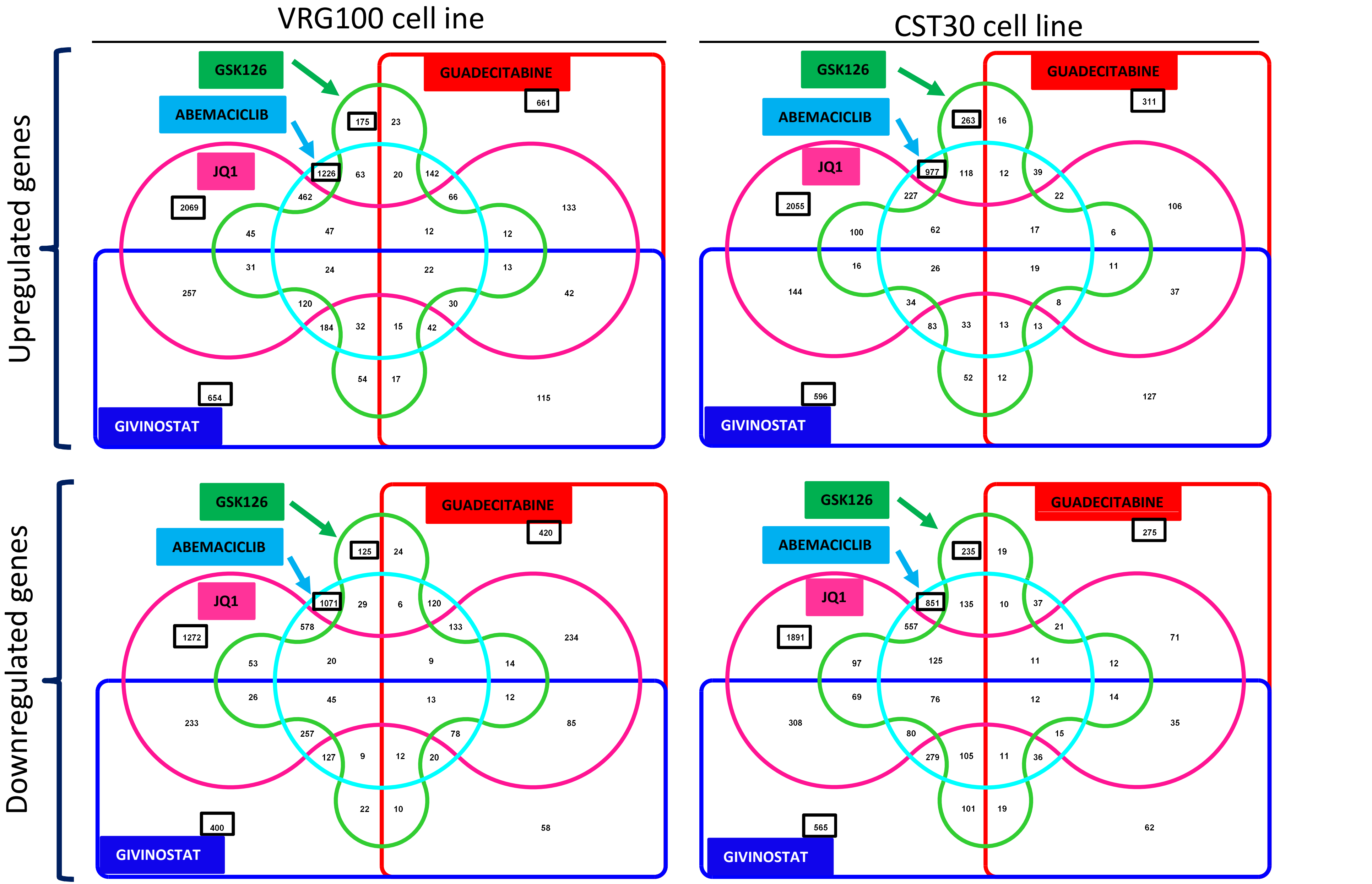


**Supplementary Figure S6**. Edwards-VENN diagram analysis of significantly modulated genes (upper panels, upregulated genes; lower panels, downregulated genes) in VRG100 (left hand panels) and CST30 (right hand panels) cell lines treated with guadecitabine (red rectangle), givinostat (blue rectangle), JQ1 (fuchsia peanut shape), GSK126 (green cogwheel) or abemaciclib (light blue circle). Numbers highlighted by a black frame represent genes modulated only by each of the drugs. All other numbers at the intersection of different colour-coded shapes represent genes co-modulated by more than one drug.

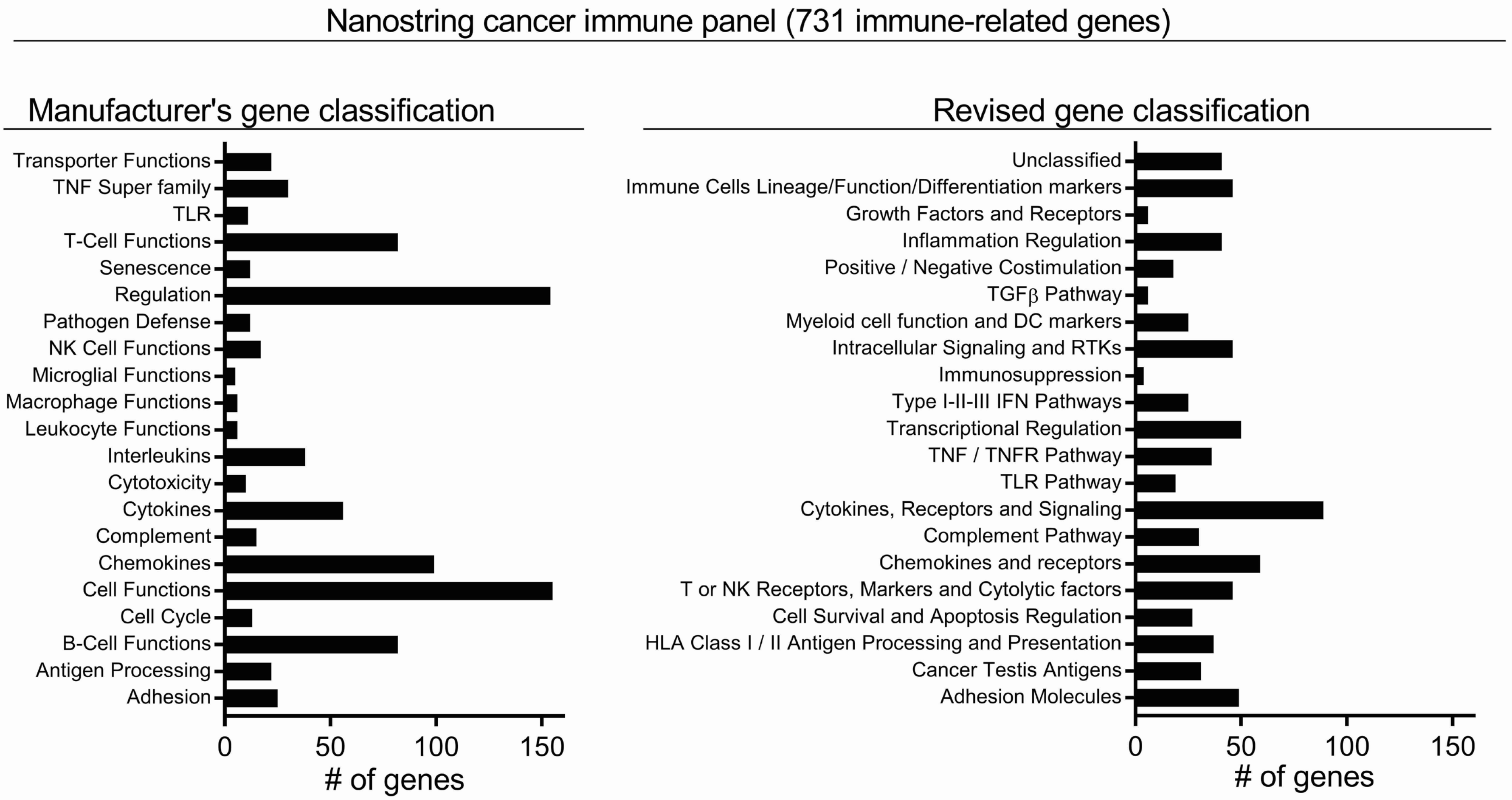


**Supplementary Figure S7**. Original (manufacturer's classification) and revised gene classification of the NanoString nCounter PanCancer Immune Profiling panel. All genes in the Nanostring panel were re-classified for function by accessing the human gene database Genecards (at http://Genecards.org) and through literature search.

Supplementary Figure S8

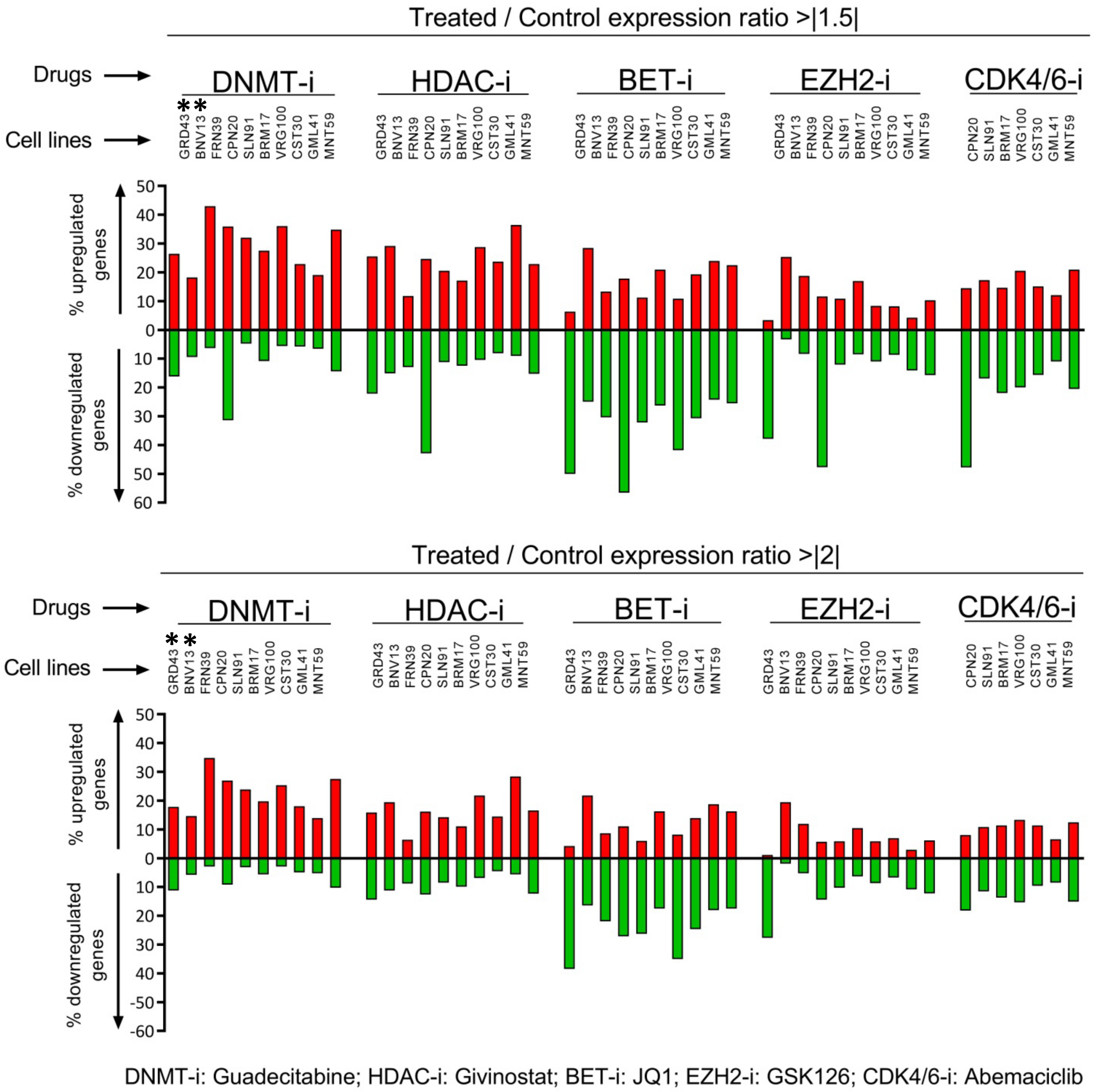


**Supplementary Figure S8**. Quantitative analysis of Nanostring data in ten melanoma cell lines treated with the indicated drugs. DNMT-i: decitabine / guadecitabine; HDAC-i:Givinostat; BET-i: JQ1; EZH2-i: GSK-126; CDK4/6 i: Abemaciclib. Upper graphs: % of genes in the Nanostring panel upregulated (red histograms) or downmodulated (green histograms) with a treated/control expression ratio >|1.5|. Lower graphs: % of genes upregulated (red histograms) or downmodulated (green histograms) with a treated/control expression ratio >|2.0|. *: these two cell lines were treated with decitabine, the active metabolite of guadecitabine.

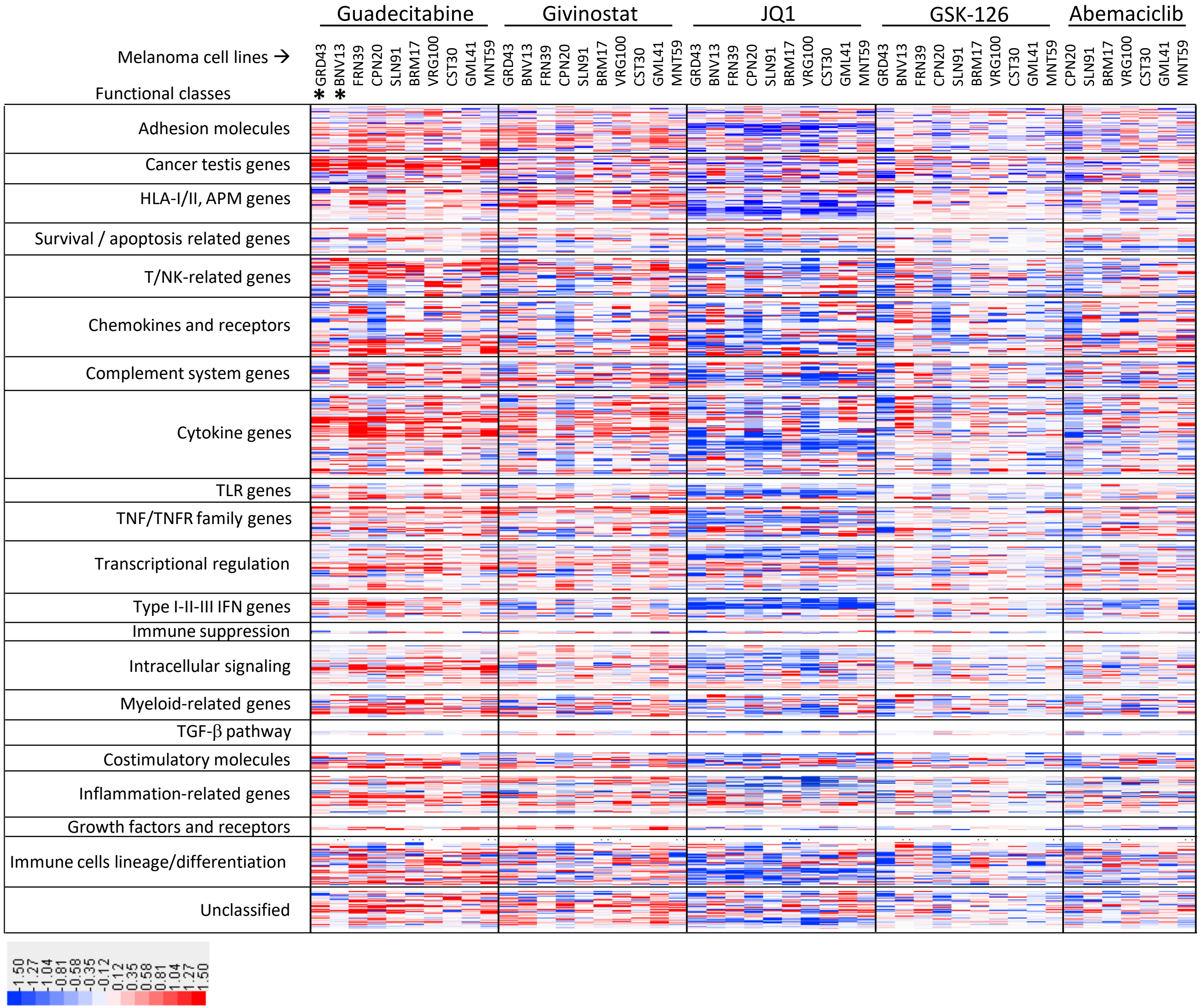


**Supplementary Figure S9. Modulation of immune-related genes in melanoma cell lines by epigenetic drugs.** Modulation of 731 genes in ten melanoma cell lines was assessed by the Nanostring Cancer Immune panel upon treatment with 4 epigenetic drugs and with the control drug Abemaciclib. Genes were clustered according to each of 21 functional classes. *: These two cell lines were treated with decitabine, the active metabolite of guadecitabine.

### Supplementary Figure S10

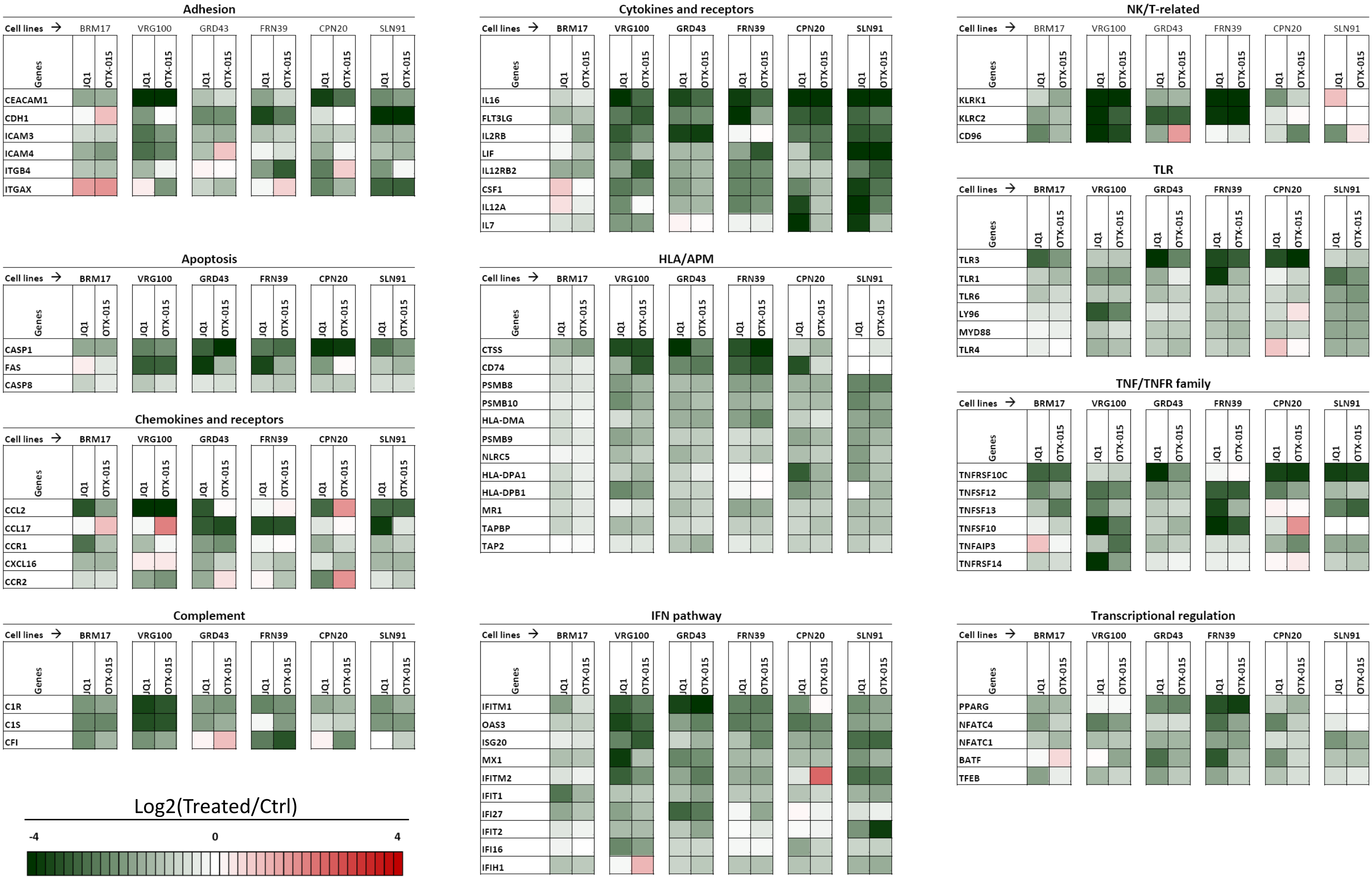


**Supplementary Figure S10**. Comparison of immune-related gene modulation by BET inhibitors JQ1 and OTX-015. Modulation of selected genes within 11 functional classes by JQ1 and OTX-015 was assessed by the Nanostring Cancer Immune Panel.

| GENE CLASS | GUADECITABINE | GIVINOSTAT | JQ1 | GSK-126 |
|---|---|---|---|---|
| Cancer testis | SSX1, MAGEB2, SSX4, CTCFL, CT45A1, CTAG1B, MAGEC2, DDX43, MAGEC1, MAGEA1, PASD1 | TMEFF2, MAGEC2, CTCFL, MAGEC1, TTK, PBK | CT45A1, BAGE, PNMA1, PBK, CTCFL | SSX1, CT45A1 |
| Class I/II Antigen Processing and Presentation | CD74, HLA-DMB, HLA-DP1, HLA-DRB4 | CD74, HLA-DRB3, HLA-DPA1, HLA-DMB, HLA-DPB1, HLA-DRB4, CD1D, HLA-DMA, HLA-DRA, MICB, HLA-B, HLA-DOB, PSMB10 | CD1C, CD1D, TAPBP, HLA-DRA, NLRC5, HLA-DMA, PSMB9, HLA-DPA1, PSMB10, CD74, PSMB8, CTSS, TAP2, HLA-B, MR1, HLA-DRB4, TAP1 | |
| Type I-II-III IFN system | IFI27, IFITM1, ISG20, IFNL2, IFITM2, IFNA7 | IFITM1, IFNA7 | IFITM1, MX1, OAS3, IFITM2, IFIT1, IFI27, ISG20, ISG15, IFIH1, IFI16, IFIT2 | IFNG |
| Chemokines and receptors | CCL20, CXCL11, XCL2, CCR1, CXCL5, CCL2, CXCL2, CXCR6, CXCR4, CXCL14 | CXCR4, CCL7, CCR6, CCL11, PPBP, CXCR3 | CXCL3, CXCL2, CX3CR1, CCR2, CXCR3, CXCL13, XCR1, CXCL9, CXCL16, CCR1, CCL2, CCL17 | PPBP, CCL11, CCR6 |
| Cytokines,receptors and signaling | IL11, IL2RG, IL24, IL13RA2, IL1B, IL2RB, IL17B, Il18, Il10, IL17RB, IL1A, IL32, CSF2, IL22RA2, IL4R, IL23A, STAT4, IL8, LIF, IL15RA, IL6, IL22RA1, IRAK2, IL1R1, IL18R1, IL1R2 | IL1B, SPP1, IL1A, Il8, IL21R, IL7R, IL10RA, IL15, IL15RA, IL3RAIL12RB1, JAK3, IL16, IL1RL1, SIGIRR | IL12RA, IL8, IL17A, IL16, FLT3LG, IL12A, LIF, IL7, CSF1, IL12RB2, IL6, SIGIRR, JAK3, IL21R, IL1R1, IL24, IRAK1, OSM, IL4R, STAT6, IL6RIL1RAP, IL13RA2, IL34, IL1RAPL2 | IL2RB, SIGIRR, IL3RA, IL1RL1, IL32 |
| TNF Super family | TNF, TNFRSF10C, TNFSF10, TNFSF11, TNFRSF1B, CD70, TNFRSF18, TNFSF15, TNFSF4, TNFSF13 | CD70, TNFRSF11A, TNFSF10,TNFRSF1B, TNFRSF18, TNFSF15, TNFSF8 | TNFRSF17, TNFRSF12A, TNFRSF8, TNFSF12, TNFRSF10C, TNFSF13, TNFSF10, TNRSF14, TNFSF15,TNFSF13BH, TNFAIP3, LTBR, TNFRSF11A, TRAF3, TNFSF11, TNFRSF13C | |
| POS-NEG costimulation | CTLA4, LAG3, ICOSLG, TIGIT, HAVCR2 | CD274, CD40, HAVCR2, CD40LG | HAVCR2, PDCD1LG2, CD200, CD40, CD80, CD274, TIGIT | LAG3, CD40LG |
| Adhesion | PECAM1, ITGB4, SELPLG, ITGB2, ITGB3, ICAM2, ITGA2B, ITGAX,THBS1, EPCAM | ITGAX, SELL, ITGA2B, ALCAM, NCAM1, FN1, ITGB3, THBS1, TH1, ITGB4 | CLEC5A, ITGB3, ALCAM, ITGA6, CD97, ITGAX, THY1, EPCAM, ICAM3, CDH1, CEACAM1 | ITGA2B, SELPLG, AMICA1 |
| Complement system | CFI, CFB, C4B, C8G, C6, MBL2, SERPING1, MASP1 | CFD, CFI, C3, SERPING1, C1R | C8B, C1R, C1S, SERPING1, CFD, C2, CFI, CD55, CFB | MASP1, C6 |
| TLR system | TLR4, TLR9, TLR2 | TLR5 | TLR10, TLR8, TLR9, TLR5, TLR3, TLR1, TLR6, LY96, MYD88, TLR4, TICAM2 | |
| intracellular signaling and RTKs | PIK3CG, INPP5D, ZAP70, AXL, SERPINB2, SOCS1, SH2D1B, TXK, LCK | SERPINB2, HCK, TXK, SH2D1B, INPP5D, LCK | SOCS1, HCK, MERTK, INPP5D, BTK, MAPK3, MAP3K5, SH2D1B, ITK, MAPK11, PIK3CD, AXL, LCK, MAP3K1 | HCK, SH2D1B, SERPINB2 |
| Transcriptional regulation | RORC, POU2F2, FOXJ1, EGR2, IRF7, PAX5, MNX1 | EGR1, EGR2, MAF, IRF5, CREB5, TCF7, FOXP3, PAX5 | MAF, EGR2, AIRE, IRF7, IRF4, PPARG, NFATC4, BATF, NFATC1, EOMES, TFEB, NFATC2, CEBPB, TPO53, CREBBP, ATF1, PAX5, FOXP3 | FOXJ1, TBX21, PAX5, AIRE |
| Inflammation regulation | SAA1, F2RL1, S100A7, NOS2A, PTGS2, CAMP, SPINK5, ANXA1, TXNIP, NOD2, ADORA2A, NLRP3, MEFV | S100A7, NOD2, F2RL1, APOE, PTGS2, TXNIP, NOS2A, PLA2G1B, MEFV, PYCARD | TXNIP, S100B, S100A7, SPINK5, IKBKE, PYCARD, CAMP, APOE, LRP1, ANP32B, ANXA1 | MEFV, PLA2G1B, ADORA2A |
| Cell survival and apoptosis regulation | ATM, BCL6, CLU | ATM, CYFIP2, BCL6, CARD11, FAS, BIRC5, CLU | ATM, ATG12, CDKN1A, BID, BCL2L1, CASP8, BIRC5, FAS, CASP1 | |
| T or NK receptors, markers and cytolytic factors | DPP4, CTSW, CD96, PRF1, SPN, CD4, KLRC2, GZMM, CD244, CD38, LCP1 | KLRK1, KLRC2, GZMB, CD96, LCP1, KIR_Inhibiting_Subgroup_1, CD244, KLRD1 | ENTPD1, CD8B, CD244, CD4, PTGDR2, DPP4, CD96, KLRK1, KLRC2 | KIR3DL2, KIR_Inhibiting_Subgroup_1, GZMH, CD8B |
| immune cells lineage/function/differ. markers | FCERIG, LY9, SLAMF1, SLAMF7, BST2, CD22, CD34, CD5, SH2D1A, SLAMF6, FCGR3A, CD79B, PTPRC, MS4A2 | CD24, RAG1, CD36, PTPRC, PRG2, CD22, LTF, FCERIG, MS4A2, FCER2 | BST2, SLAMF7, FCER1G, CD36, LTF, ADA, FCGR2A, BST1, RAG1, CD22, LILRB2, MME, FCER2, CD83, PLA2G6, MS4A2, FCER1A | CD5, FCER2, FCER1G, IGLL1, FCGR3A, FCER1A |
| Myeloid cell function and DC markers | CD14, TREM2, MST1R, THBD, NRP1, SLC11A1, CD207, CHIT1 | CD14, TREM2, NRP1, CHIT1 | CHIT1, NRP1, TREM2 | MST1R, NCF4 |
| NOT classified | LCN2, LAMP3, LRRN3, DMBT1, TPSAB1, PMCH, PLAU, PLAUR, SMPD3, HSD11B1, FPR2,FUT7, MUC1 | NEFL, ABCB1, LAMP3, FEZ1, DMBT1, PLAUR, F12, MUC1 | FPR2, NEFL, ABCB1, LAMP2, MUC1, F12, CTSH, CTSG, CTSL, SYT17, TPSAB1, PLAUR, DMBT1 | F13A1 |

**Supplementary Figure S11**. **Immune-related signature of epigenetic drugs in melanoma**. The table shows the genes observed upregulated (red) or downmodulated (green) with the same direction of change in at least 6/10 cell lines and showing a Treated/Ctrl ratio >|1.5|.

Supplementary Figure S12.

A

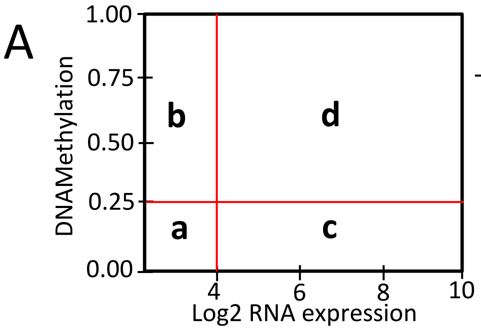


| Class | Log2 RNA Expression | DNA Methylation | Meaning |
|---|---|---|---|
| a | <4 | <0.25 | Expression^LOW^/methylation^LOW^ |
| b | <4 | >0.25 | Expression^LOW^/methylation^HIGH^ |
| c | >4 | <0.25 | Expression^HIGH^/methylation^LOW^ |
| d | >4 | >0.25 | Expression^HIGH^/methylation^HIGH^ |

C

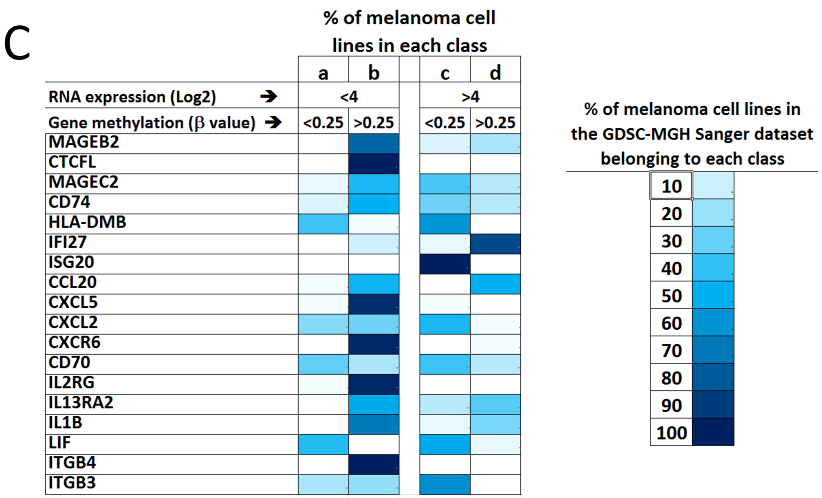


B

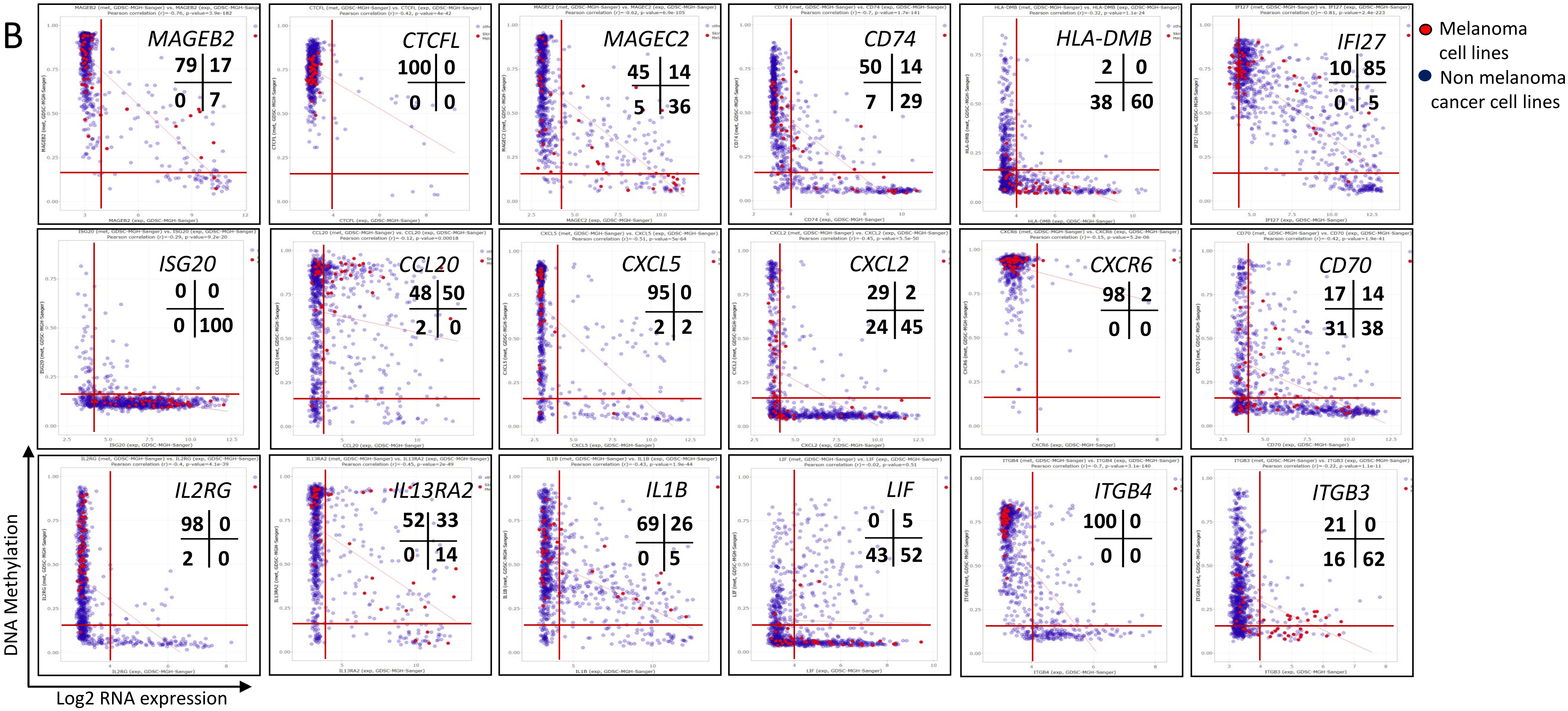

**Supplementary Figure S12. Expression/methylation relationship, for selected genes in the guadecitabine-specific gene signature, among melanoma cell lines present in the GDSC-MGH Sanger database. A.** The expression/methylation relationship for genes of interest, based on data retrieved from the CellMiner CDB web application, were visualized in scatter plots where four main quadrants (a,b,c,d, each characterized by distinct levels of expression and DNA methylation) could be defined by the indicated thresholds. **B**. Representative scatter plots for 18 genes in the guadecitabine signature. Red dots indicate single melanoma cells lines, blue dots represent cell lines of other histological origin. Numbers in each gene-specific scatter plot represent the % of all melanoma cell lines classified in one of the four expression/methylation quadrants. **C.** The data in panel B were subsequently summarized in a color-coded tabular form where the percentage of cell lines in each of the four expression/methylation quadrants is shown by shades of blue color.

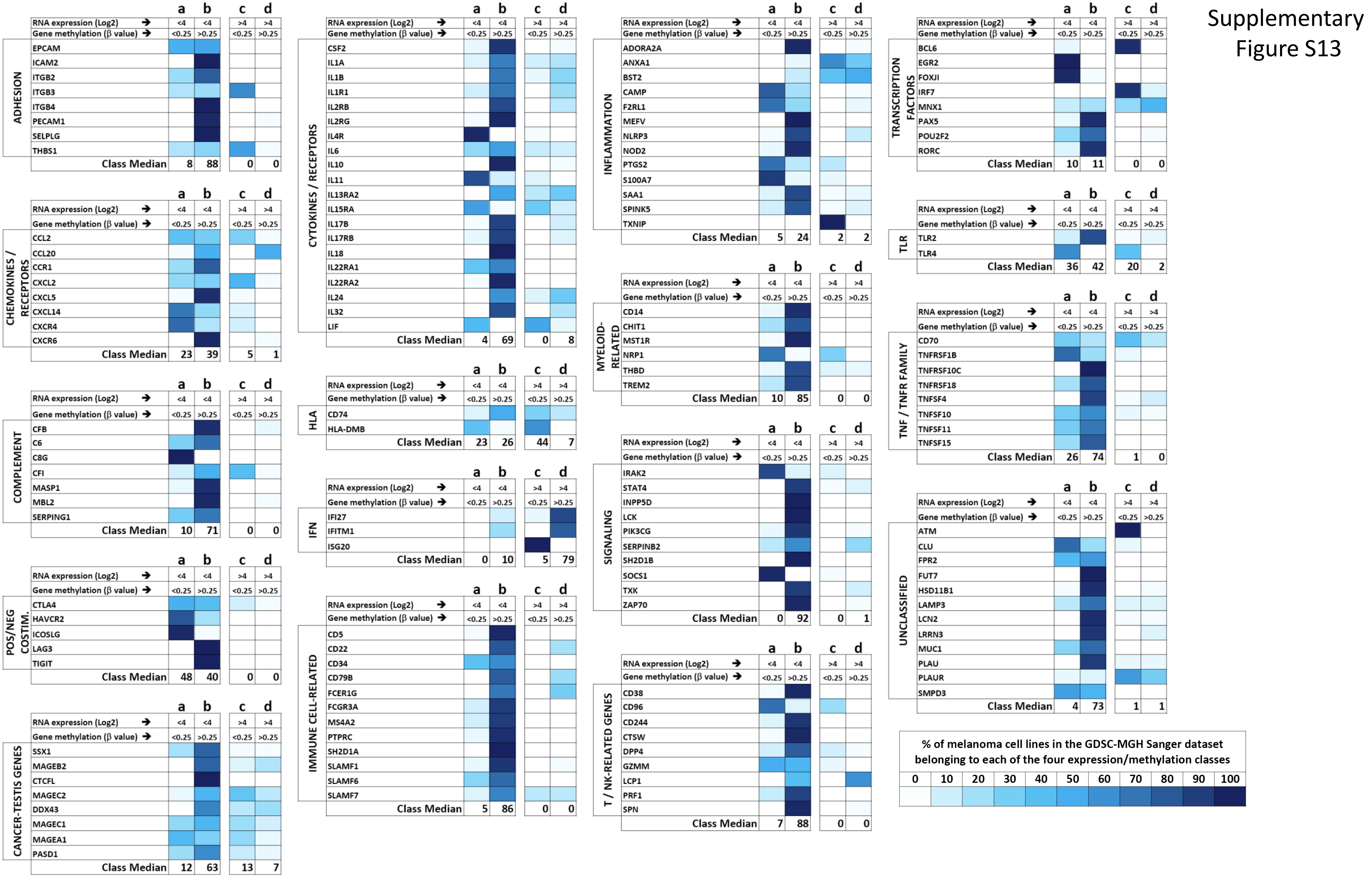


**Supplementary Figure S13.** Expression/methylation relationship of genes belonging to the guadecitabine-specific signature according to the GDSC-MGH Sanger melanoma dataset. For each gene of interest, the % of cell lines in the dataset [classified in each of 4 expression/methylation groups (a,b,c,d) according to the criteria defined in Supplementary Fig. S12], was visualized by the color code indicated at the bottom of the figure.

Supplementary Figure S14

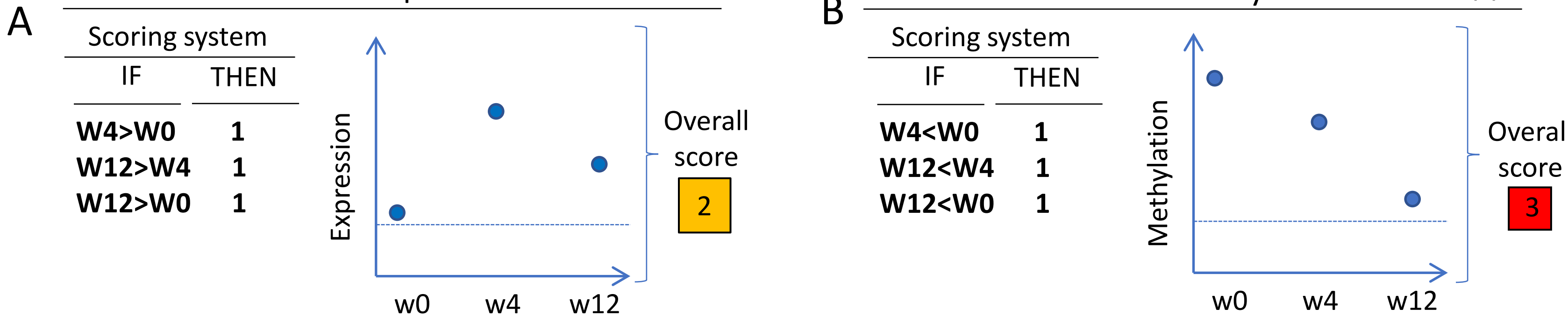


C

Responding Patients

| Gene symbol | PT#3 EXPR | PT#3 METH | | PT#4EXPR | PT#4 METH | | PT#5 EXPR | PT#5 METH | |
|---|---|---|---|---|---|---|---|---|---|
| MAGEB2 | 1 | 3 | • | 2 | | | 1 | 2 | • |
| CTCFL | 1 | 3 | • | 3 | 0 | | 0 | 1 | |
| CT45A1 | 0 | 2 | | 0 | 1 | | 0 | 2 | |
| MAGEC2 | 1 | 1 | • | 2 | 0 | | 1 | 0 | |
| MAGEC1 | 1 | 1 | • | 3 | 1 | • | 1 | 0 | |
| MAGEA1 | 1 | 2 | • | 2 | 3 | • | 0 | 0 | |
| PASD1 | 2 | 2 | • | 2 | 1 | • | 2 | 0 | |
| MAGEA3 | 1 | 1 | • | 0 | 1 | | 1 | 1 | • |
| TMEFF2 | 0 | 1 | | 0 | 1 | | 2 | 3 | • |
| GAGE1 | 1 | 3 | • | 3 | 1 | • | 0 | 0 | |
| PNMA1 | 3 | 2 | • | 1 | 2 | • | 1 | 0 | |
| ROPN1 | 1 | 3 | • | 2 | 1 | • | 2 | 2 | • |
| CTAGE1 | 1 | 1 | • | 1 | 2 | • | 0 | 0 | |
| SPA17 | 1 | 1 | • | 2 | 2 | • | 2 | 0 | |
| MAGEA4 | 3 | 2 | • | 2 | | | 1 | 0 | |
| SYCP1 | 0 | 2 | | 1 | 1 | • | 2 | 3 | • |
| SPANXB1 | 0 | 1 | | 0 | 0 | | 2 | 2 | • |
| SPO11 | 0 | 1 | | 1 | | | 0 | 0 | |
| TTK | 1 | 0 | | 0 | | | 2 | 0 | |
| PBK | 0 | 2 | | 2 | 1 | • | 2 | 2 | • |
| PRM1 | 0 | 2 | | 0 | | | 0 | 2 | |

Non Responding patients

| Gene symbol | PT#1 EXPR | PT#1 METH | | PT#2EXPR | PT#2METH | | PT#6 EXPR | PT#6 METH | | PT#7 EXPR | P#T7 METH | | PT#8 EXPR | PT#8 METH | |
|---|---|---|---|---|---|---|---|---|---|---|---|---|---|---|---|
| MAGEB2 | 1 | 1 | • | 0 | 1 | | 1 | | | 2 | 3 | • | 1 | 1 | • |
| CTCFL | 0 | 3 | | 1 | 1 | • | 1 | 0 | | 0 | 0 | | 2 | 1 | • |
| CT45A1 | 0 | 3 | | 1 | 2 | • | 1 | 1 | • | 0 | 0 | | 0 | 1 | |
| MAGEC2 | 1 | 2 | • | 1 | 1 | • | 1 | 1 | • | 2 | 1 | • | 3 | 0 | |
| MAGEC1 | 2 | 1 | • | 1 | 0 | | 0 | 1 | | 1 | 1 | • | 2 | | |
| MAGEA1 | 1 | 2 | • | 0 | 1 | | 0 | 1 | | 3 | 0 | | 1 | 0 | |
| PASD1 | 3 | 1 | • | 1 | 1 | • | 1 | 0 | | 0 | 3 | | 0 | 3 | |
| MAGEA3 | 3 | 3 | • | 0 | 1 | | 0 | 1 | | 2 | 2 | • | 2 | 3 | • |
| TMEFF2 | 2 | 0 | | 0 | 1 | | 1 | 1 | • | 1 | 2 | • | 0 | 2 | |
| GAGE1 | 0 | 3 | | 1 | 1 | • | 1 | 0 | | 1 | 3 | • | 0 | 1 | |
| PNMA1 | 2 | 3 | • | 3 | 1 | • | 0 | 0 | | 3 | 1 | • | 0 | 2 | |
| ROPN1 | 2 | 0 | | 0 | 1 | | 1 | 0 | | 1 | 3 | • | 2 | 0 | |
| CTAGE1 | 2 | | | 2 | 0 | | 0 | 0 | | 2 | 0 | | 1 | 0 | |
| SPA17 | 2 | 1 | • | 2 | 1 | • | 1 | 0 | | 2 | 2 | • | 1 | 2 | • |
| MAGEA4 | 1 | | | 0 | 2 | | 0 | | | 3 | | | 2 | | |
| SYCP1 | 3 | 3 | • | 3 | 0 | | 0 | 1 | | 0 | 1 | | 0 | 2 | |
| SPANXB1 | 2 | 3 | • | 1 | 0 | | 1 | | | 0 | 1 | | 0 | 1 | |
| SPO11 | 0 | | | 0 | 0 | | 0 | | | 0 | | | 2 | | |
| TTK | 2 | | | 3 | 0 | | 1 | | | 2 | | | 1 | 0 | |
| PBK | 2 | 1 | • | 0 | 3 | | 0 | 1 | | 2 | 3 | • | 0 | 1 | |
| PRM1 | 0 | 0 | | 0 | 1 | | 0 | | | 0 | | | 0 | | |

**Supplementary Figure S14. Changes in gene expression and gene methylation for the cancer testis class of genes in tumor biopsies from melanoma patients enrolled in the NIBIT-M4 trial. A, B**. Outline of the scoring system adopted for potentially observable changes. To simplify data analysis, only instances of increase in gene expression and decrease in gene-specific methylation were considered. By such criteria, an increase in gene expression at any of the three possible comparisons (w4 vs w0, w12 vs w4 and w12 vs w0) receives a score = 1; in the instance of the example shown in A the overall score for gene expression change during therapy is=2. In the instance of the example shown in B the overall score for gene methylation change during therapy is = 3. **C**. Observed changes in expression/methylation of cancer testis genes in NIBIT-M4 neoplastic tissues, classified according to the score system defined in panels A and B. Black dots highlight genes showing both increase of expression and decrease of methylation in some of the three possible comparisons (w4 vs w0, w12 vs w4 and w12 vs w0). w0: baseline tumor sample; w4: on-treatment biopsy at week 4; w12: on treatment biopsy at week 12.

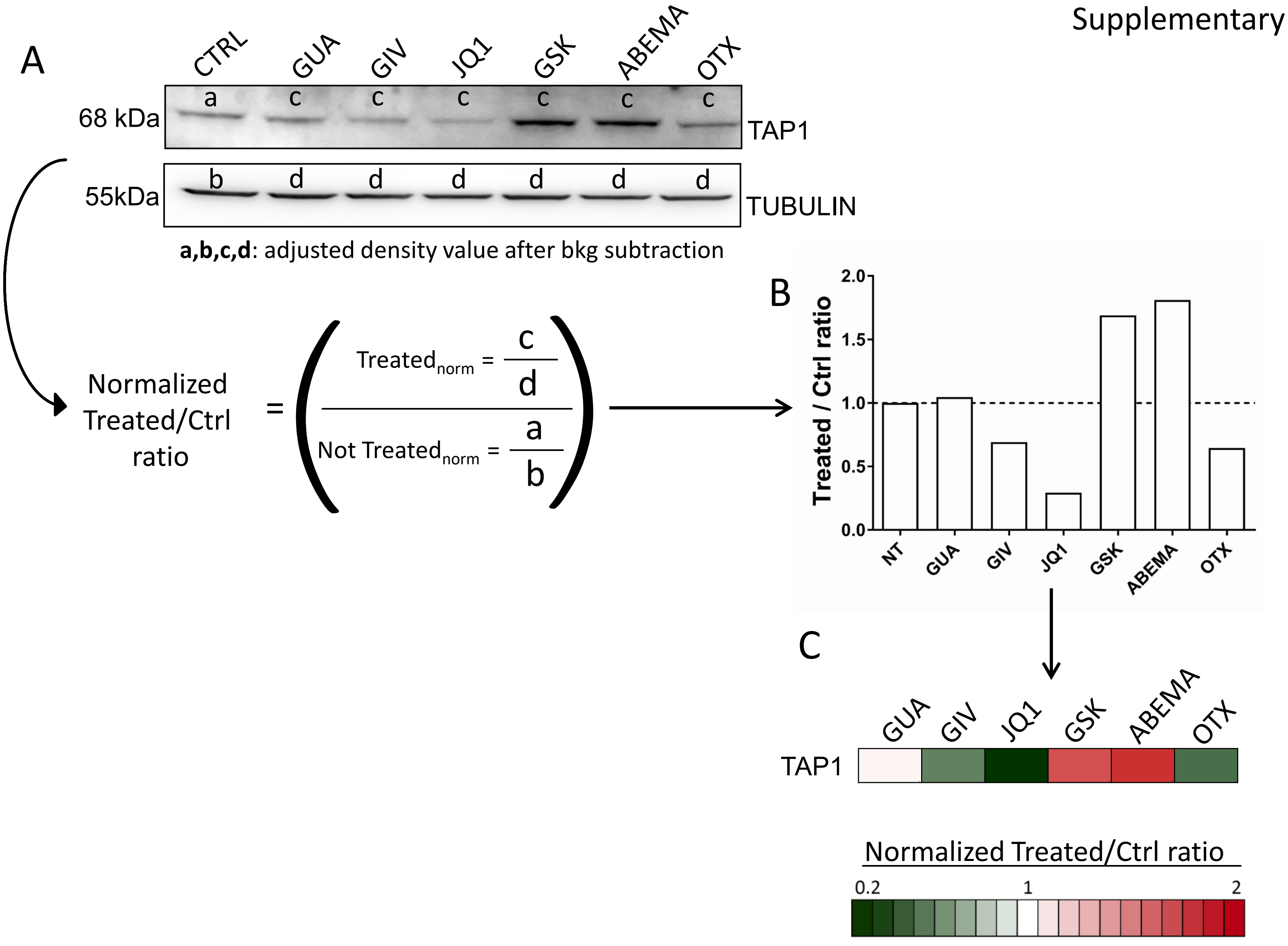


**Supplementary Figure S15**. **Outline of the strategy for quantitative analysis and visualization of western blot data. A.** Normalized treated/control ratios were computed on the basis of background-adjusted density values. **B,C**. Treated/control ratio values were then converted to a color-coded strip allowing direct visualization of the effect of each drug on markers of interest. CTRL. Untreated cells; GUA: guadecitabine; GIV: givinostat; GSK: GSK126; ABEMA: abemaciclib; OTX:OTX-015.

Supplementary Figure S16

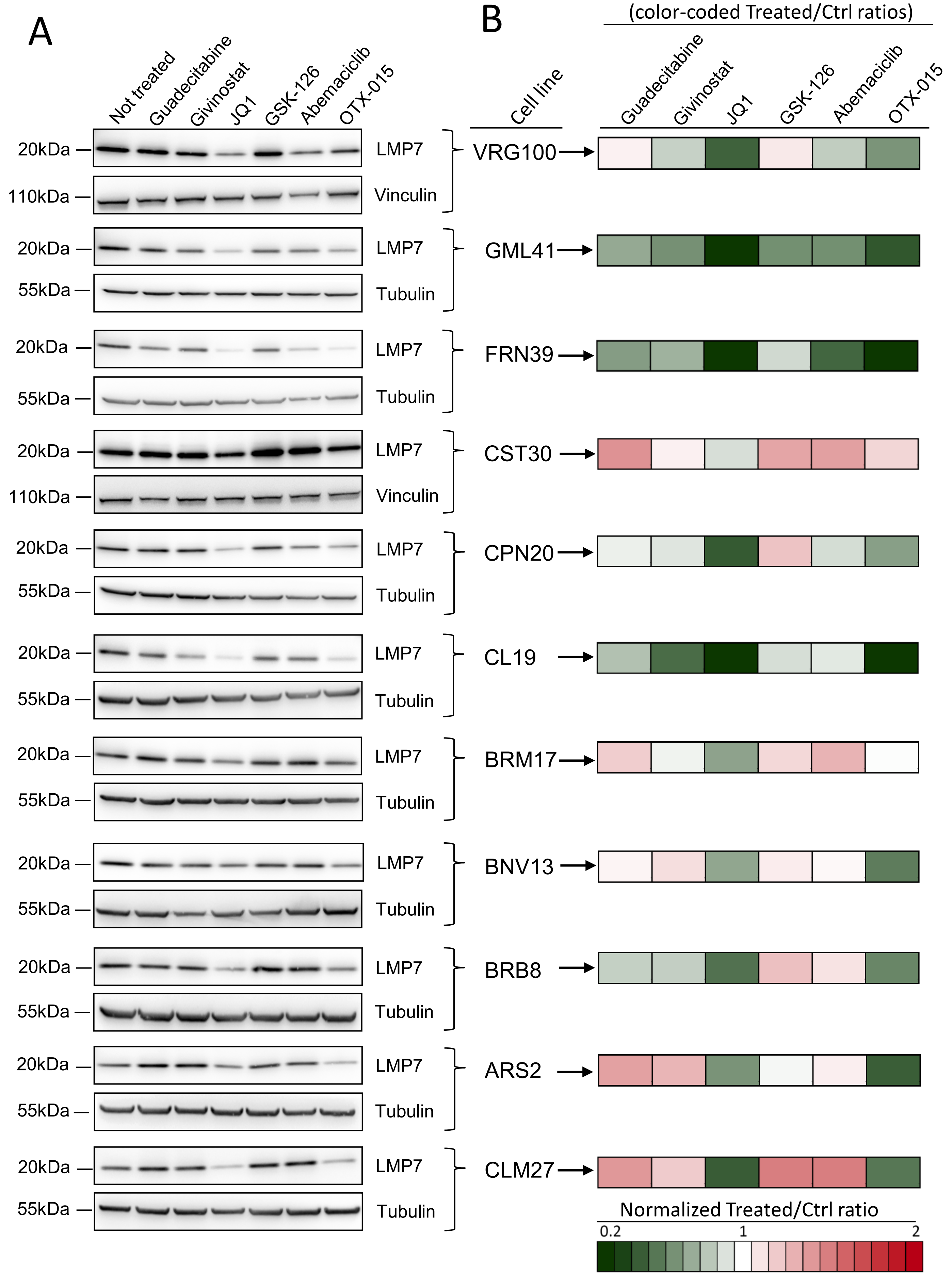


**Supplementary Figure S16**. Quantitative western blot analysis and visualization of the modulation of LMP7 by epigenetic drugs in 11 melanoma cell lines. **A**. Original western blot images. **B**. Color-coded normalized treated/control ratios as defined in Fig. S15.

Supplementary Figure S17.

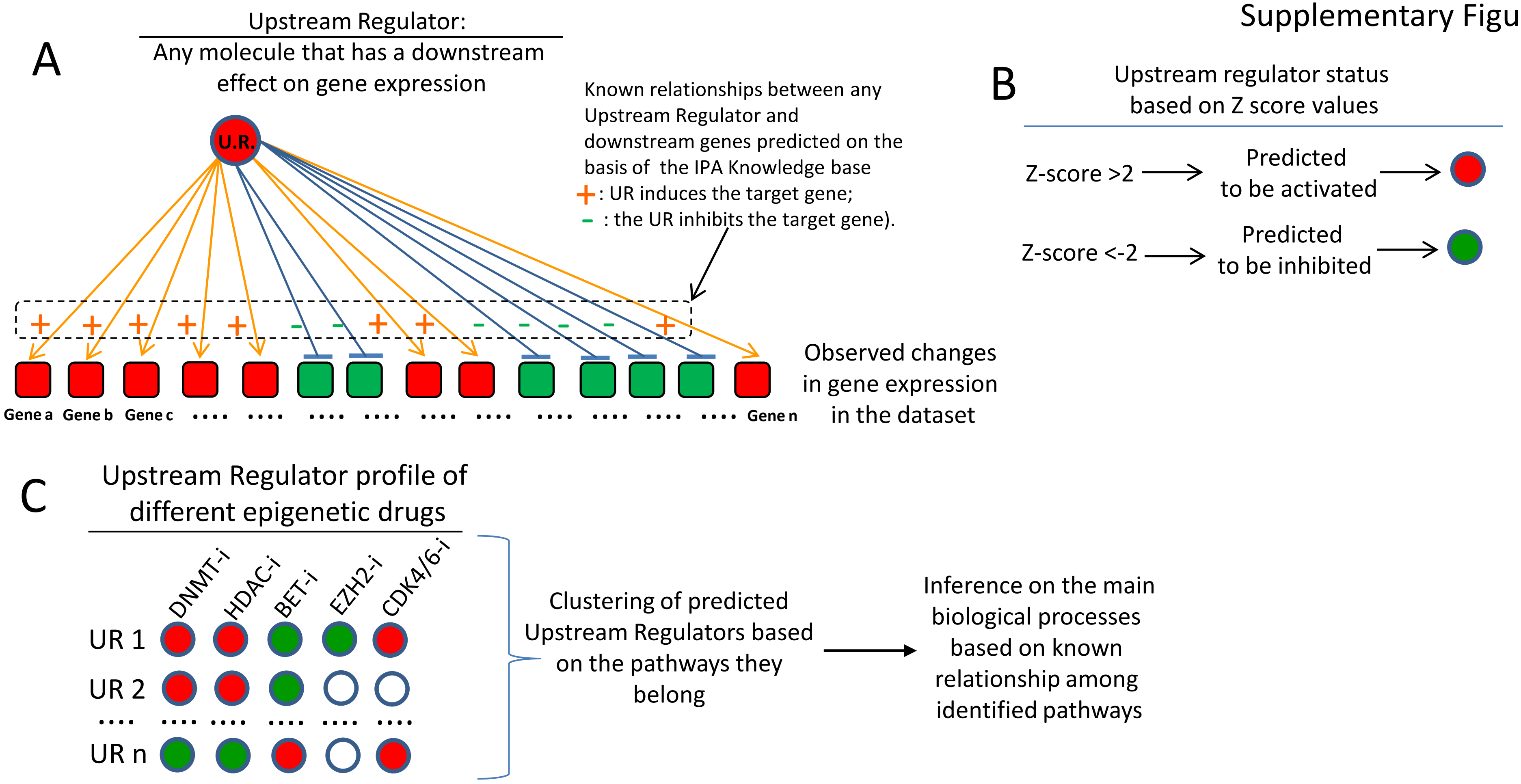


**Supplementary Figure S17**. **Pipeline of data analysis based on Upstream Regulators (UR) identified by IPA**. **A**. In this schematic, squares represent genes, while circles represent URs. Circle color denotes predicted UR status (red: activation; green: inhibition). Square color denotes observed gene expression change (red: upregulation; green: downmodulation). An UR is any molecule that can have a downstream effect on gene expression. The IPA knowledge base, built into the application, identifies the relationships between any set of genes being observed as significantly modulated in the dataset and the UR that controls them (relationship measured through a P value of overlap between a set of genes and any given upstream regulator). **B**. Depending on the type of relationship between the set of genes and an UR, and on the observed changes in gene expression, IPA computes a Z score statistics whose meaning is to infer the activation status ("activated" or "inhibited") of the UR. Only Z scores greater than 2 or smaller than -2 were considered significant. **C**. For each drug, the overall UR profile can be identified in terms of identity of the molecules (UR1, UR2…URn) and of their predicted activation status. Different URs can then be clustered together based on the common biological pathways they belong. Finally, an inference can be made on the biological processes being modulated by each drug based on known relationships between the identified pathways.

Supplementary Figure S18

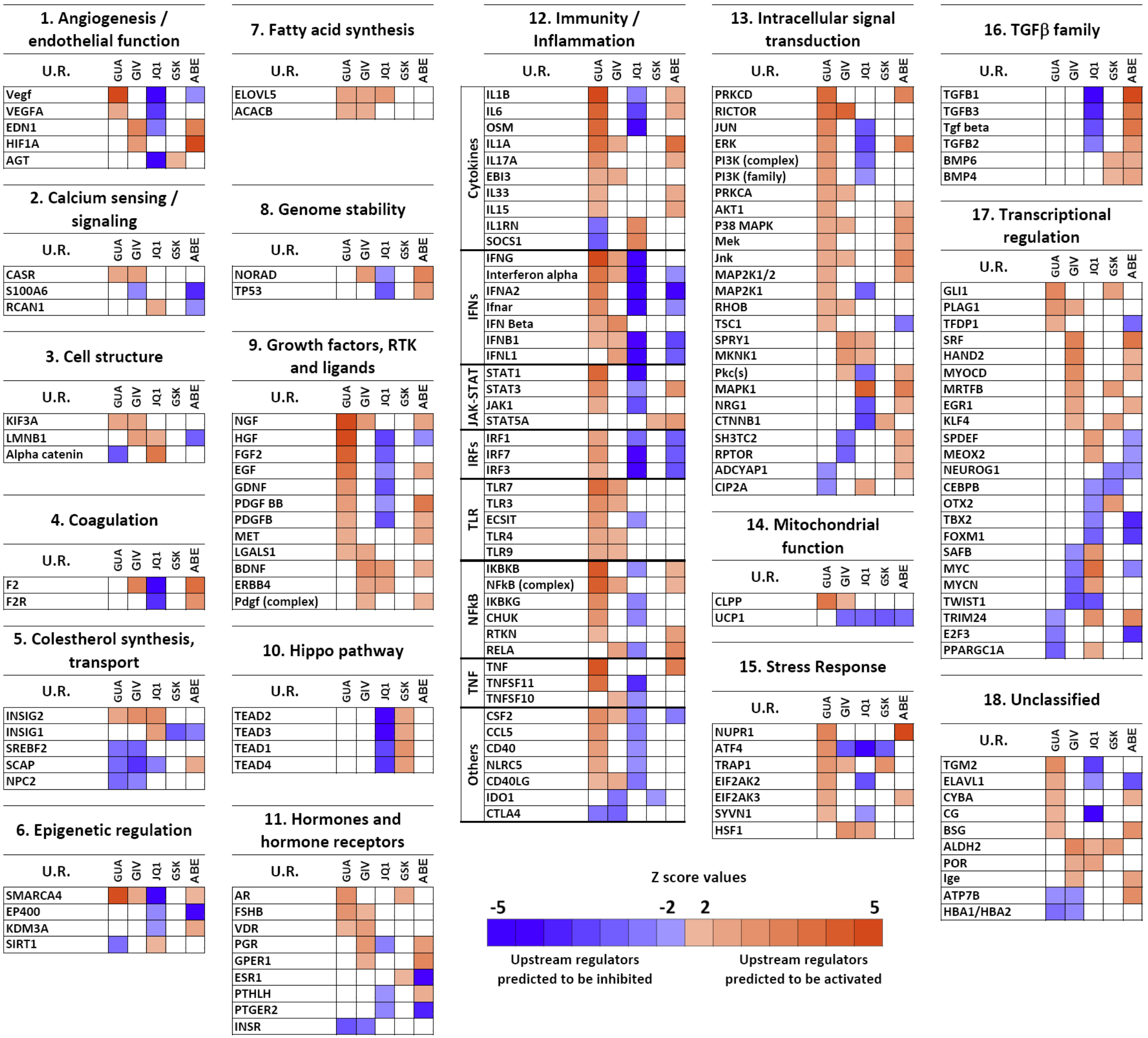


**Supplementary Fig. S18. Classification of URs significantly modulated by at least two different drugs in melanoma cell line VRG100.** URs significantly modulated by at least two different drugs were grouped into 18 functional classes. Each UR was selected based on a significant Z score (>|2| and a significant p value for association with specific sets of modulated genes by each drug. Z score values of each UR are shown by a color code indicating prediction of UR inhibition (blue) or prediction of UR activation (red). GUA: guadecitabine, GIV: givinostat, GSK: GSK-126; ABE: abemaciclib.

Supplementary Figure S19

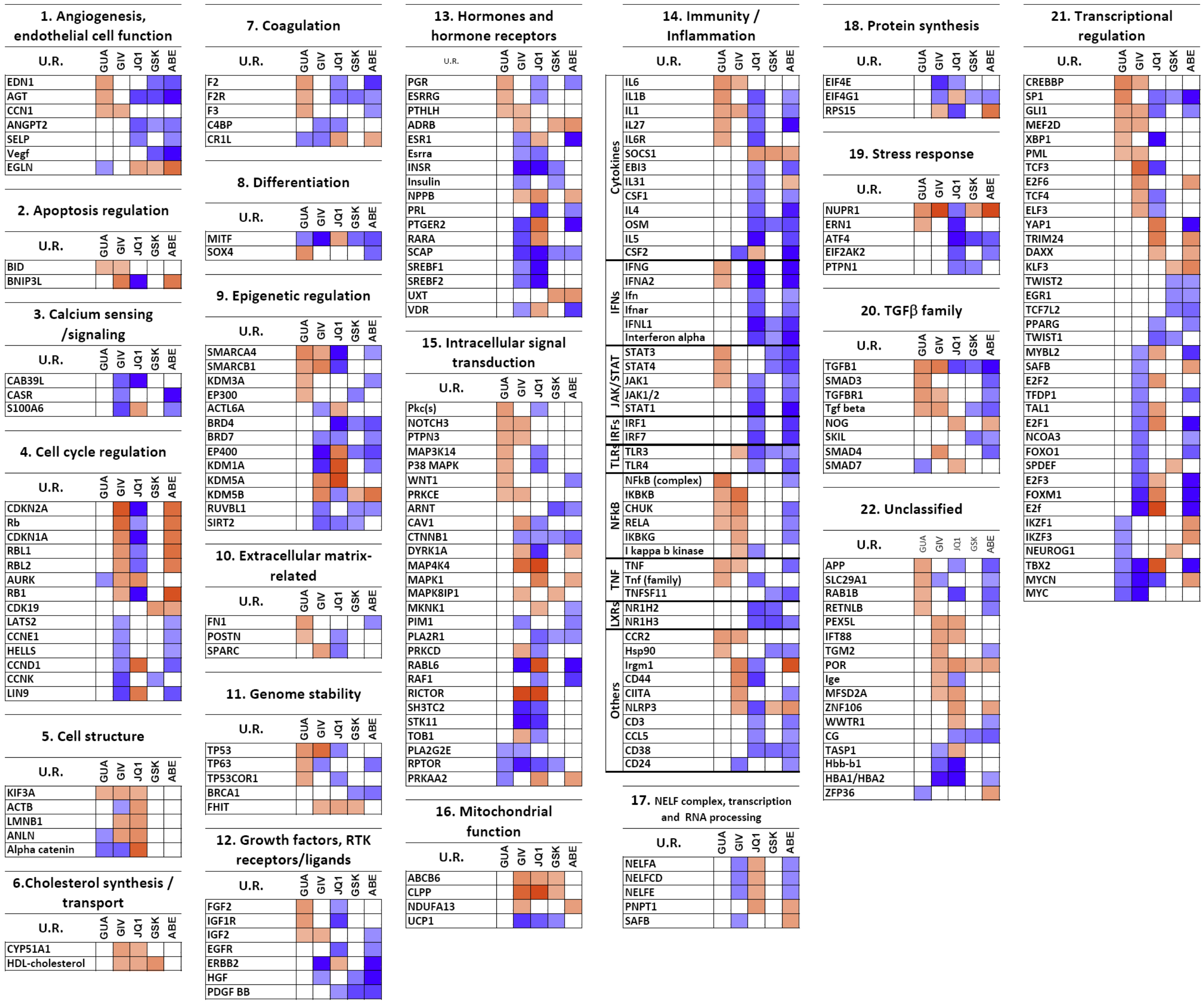


**Supplementary Fig. S19. Classification of URs significantly modulated by at least two different drugs in melanoma cell line CST30.** URs significantly modulated by at least two different drugs were grouped into 18 functional classes. Each UR was selected based on a significant Z score (>|2| and a significant p value for association with specific sets of modulated genes by each drug. Z score values are shown by a color code indicating prediction of UR inhibition (blue) or prediction of UR activation (red). GUA: guadecitabine, GIV: givinostat, GSK: GSK-126; ABE: abemaciclib.

#### Supplementary Figure S20

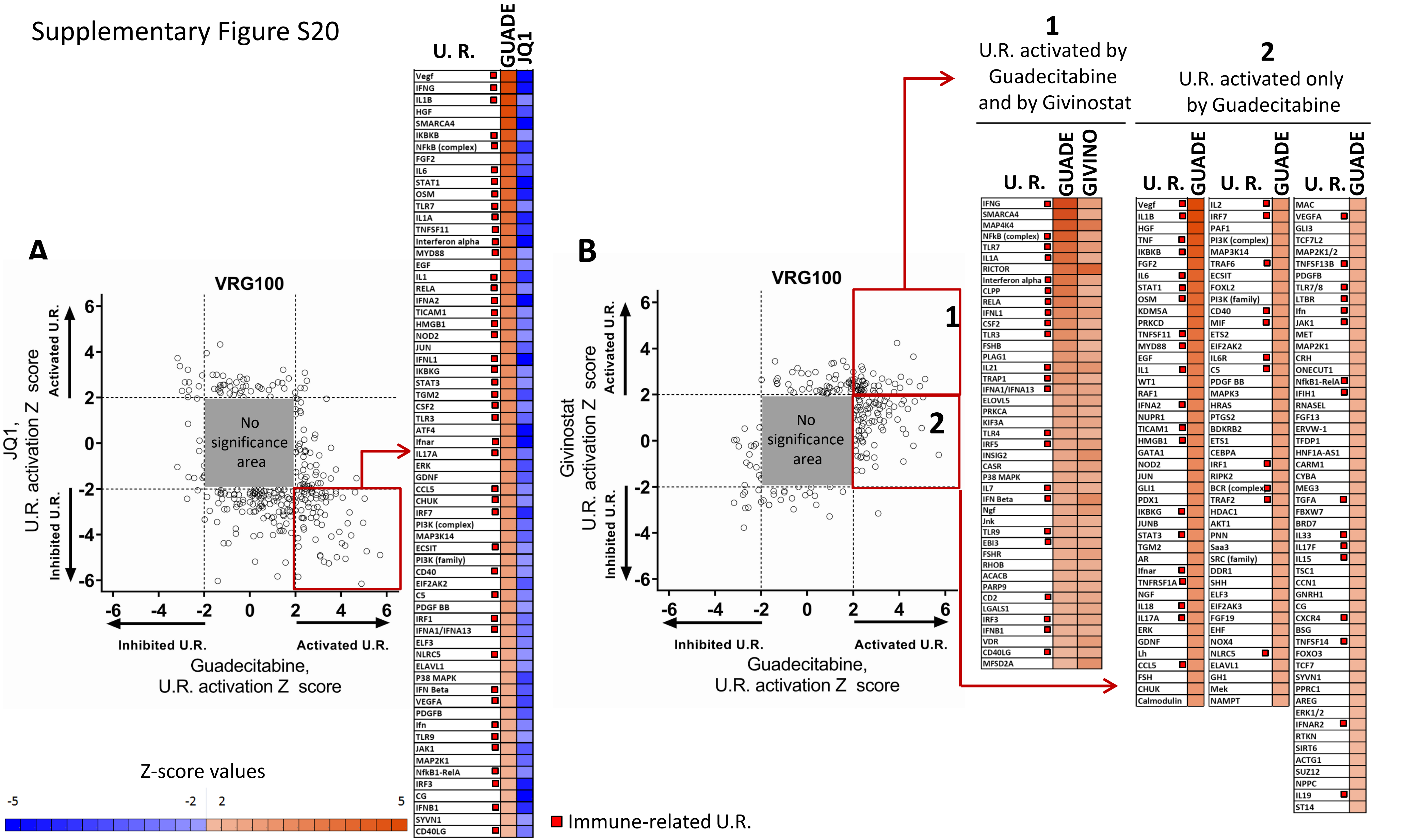


**Supplementary Figure S20. Comparison of all URs predicted to be activated or inhibited by Guadecitabine vs JQ1 and by Guadecitabine vs Givinostat**. **A**. Scatter plot of significantly modulated URs by Guadecitabine and JQ1. The grey square represents the area of no-significance, i.e. URs with Z<|2| and p value >0.05. URs in the square highligthed in red, representing factors showing opposite type of predicted modulation (activation by guadecitabine and inhibition by JQ1), are listed in the table on right of the scatter plot. **B**. Scatter plot of significantly modulated URs by guadecitabine and givinostat. URs in square 1, highligthed in red, represent factors showing predicted activation by both guadecitabine and givinostat; URs in square 2 represent factors predicted to be activated only by guadecitabine and not by givinostat. URs in square 1 and 2 are listed in the tables on the right side of the scater plot. Red squares identify immune-related Urs.

A

| | | Shared upstream regulators activated by Guadecitabine in-vivo and in-vitro | |
|---|---|---|---|
| | | In vivo | In vitro |
| Pathway | Upstream Regulator | Z score values, tumor nodules from treated vs control mice, cell line 195 | Average Z score values, treated vs untreated melanoma cell lines |
| green | IFNG | 7.05 | 4.20 |
| green | IFNA2 | 6.83 | 3.24 |
| green | IFNL1 | 6.28 | 3.14 |
| green | IRF1 | 5.26 | 2.78 |
| green | Ifnar | 5.06 | 3.05 |
| green | STAT1 | 4.97 | 3.68 |
| green | EIF2AK2 | 4.88 | 2.85 |
| | SMARCA4 | 4.70 | 2.75 |
| green | IRF3 | 4.40 | 2.85 |
| blue | TGM2 | 4.08 | 2.89 |
| | DOCK8 | 3.30 | 3.03 |
| red | TNF | 3.26 | 4.60 |
| red | TICAM1 | 3.19 | 3.56 |
| red | IL1B | 3.15 | 4.06 |
| red | SASH1 | 3.13 | 3.07 |
| green | SAMSN1 | 2.84 | 3.15 |
| blue | RELA | 2.56 | 3.37 |
| blue | IKBKB | 2.53 | 3.30 |
| green | DDX58 | 2.53 | 2.51 |
| red | TLR7 | 2.50 | 3.71 |
| red | TLR3 | 2.30 | 3.84 |
| blue | CD40LG | 2.23 | 3.00 |
| | ARHGAP21 | 2.11 | 2.59 |

| Code | Pathway |
|---|---|
| green | Type-I/III IFN |
| blue | NF-kB |
| red | TLR |

#### Canonical pathway “Interferon signaling”

B

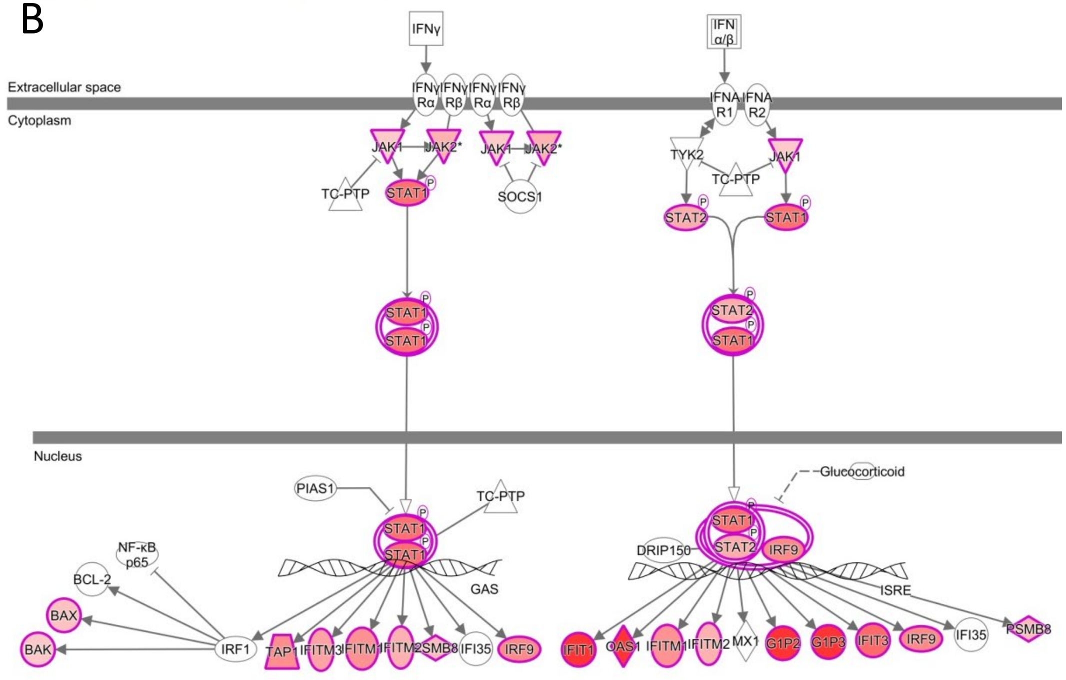


**Supplementary Figure S21. Comparison of URs activated by guadecitabine in-vitro and in-vivo. A.** Table of top URs activated by guadecitabine in tumor nodules from mice bearing a human melanoma xenograft (cell line 195) and treated with this drug. Z score values computed from gene expression data of treated mice vs control mice (first column) are compared, for each UR, to the average Z score value observed in vitro in melanoma cell lines treated with guadecitabine. **B**. Canonical pathway analysis of IFN-γ and IFN-α/β pathways modulated by guadecitabine in vivo in tumor nodules from treated vs control mice. Genes highlighted in red were observed as significantly upregulated by Guadecitabine.
